# Intracellular Retention of Estradiol is Mediated by GRAM Domain-Containing Protein ASTER-B in Breast Cancer Cells

**DOI:** 10.1101/2024.05.16.594581

**Authors:** Hyung Bum Kim, W. Lee Kraus

## Abstract

Elevated blood levels of estrogens have been associated with poor prognosis in estrogen receptor-positive (ER+) breast cancers, but the relationship between circulating hormone levels in the blood and intracellular hormone concentrations are not well characterized. We observed that MCF-7 cells treated acutely with 17β-estradiol (E2) retain a substantial amount of the hormone even upon removal of the hormone from the culture medium. Moreover, global patterns of E2-dependent gene expression are sustained for hours after acute E2 treatment and hormone removal. While circulating E2 is sequestered by sex hormone binding globulin (SHBG), the potential mechanisms of intracellular E2 retention are poorly understood. We found that a mislocalization of a steroid-binding GRAM-domain containing protein, ASTER-B, to the nucleus, which is observed in a subset of breast cancer patients, is associated with higher cellular E2 retention. Accumulation and retention of E2 are related to the steroidal properties of E2, and require nuclear localization and steroid binding by ASTER-B, as shown using a panel of mutant ASTER-B proteins. Finally, we observed that nuclear ASTER-B-mediated E2 retention is required for sustained hormone-induced ERα chromatin occupancy at enhancers and gene expression, as well as subsequent cell growth responses. Our results add intracellular hormone retention as a mechanism controlling E2-dependent gene expression and downstream biological outcomes.

**Significance:** This study demonstrates how E2 can be accumulated and retained intracellularly to drive a pro-proliferative gene expression program in ER+ breast cancer cells. Mechanistically, intracellular E2 retention is mediated in part by mislocalized nuclear ASTER-B, which binds estradiol to support the functions of ER, including E2-mediated gene expression

## Introduction

Estrogens are a class of steroid hormones required for a diverse range of physiological and pathological processes. The most abundant form, 17β-estradiol (E2), acts as the primary mitogen for estrogen receptor positive (ER+) breast cancers, which make up 80% of breast cancer molecular subtypes (1). Higher circulating concentrations of E2 are associated with a higher risk of breast cancer and poorer patient outcomes (2–4). Physiological levels of estrogens are regulated by their production, degradation, and sequestration (5). Biological and pharmacological methods of reducing E2 levels in breast cancers have generally resulted in better prognosis. For example, inhibitors such as anastrozole, exemestane, or letrozole which target aromatases, key enzymes responsible for E2 production, effectively reduce circulating E2 concentrations, halting ER+ breast cancer growth and recurrence (6–9). In addition to synthesis, circulating estrogens are bound by sex hormone binding globulin (SHBG), which binds and sequester steroid hormones, effectively limiting their bioavailability (10).

While the use of aromatase inhibitors has proven effective in reducing circulating E2 concentrations, intratumoral E2 concentrations are often elevated (11,12). Intratumoral concentrations of E2 have been found to be upwards of 10-fold higher than circulation concentrations (13). These elevated local concentrations have been shown to facilitate the growth of ER+ breast cancers despite reduced circulating E2. While aberrant intratumoral hormone production has been characterized, the role of cellular accumulation of E2 is still poorly understood (14,15). Therefore, mechanistic insights into E2 accumulation in breast cancer cells would be invaluable for better understanding effective E2 concentrations in ER+ breast cancers.

Recently, ASTER-B has been implicated in E2 homeostasis. ASTER-B (encoded by *GRAMD1B*) is a GRAM-domain containing protein involved in nonvesicular cholesterol transport between the endoplasmic reticulum and plasma membrane. The protein is normally localized to the endoplasmic reticulum and has recently been shown to play a role in pathways involving estrogen production in the ovaries (16). Additionally, biochemical studies examining the binding of a broad range of cholesterol-derived molecules show that E2 can bind directly to ASTER-B, albeit with weak affinity (17). While these studies have begun to draw connections between ASTER-B and estrogen levels, the role of ASTER-B in ER+ breast cancer pathology has not been explored.

Here we characterize a role for ASTER-B in the intercellular accumulation and retention of E2 contributing to E2-regulated gene regulation and mitogenic effects.

## Results

### Breast cancer cells retain intracellular E2 and sustain gene expression after acute E2 treatment followed by hormone washout

In order to determine the level of E2 cellular retention in ER+ breast cancer cells, MCF-7 cells were acutely treated (40 min) with 100 nM E2 followed by incubation in E2-free medium (40 min) before measuring cellular E2 content (Fig. 1A). Breast cancer cells lines expressing high ERα expression (MCF-7), moderate ERα expression (T-47D), and no ERα expression (MDA-MB-231) were tested. None of the cell lines had substantial E2 content prior to treatment. Both ER+ breast cancer cells lines accumulated a substantial level of E2 at a concentration similar to that of the treatment concentration (i.e., 100 nM). Moreover, MCF-7 cells retained almost half the treatment concentration of E2 after washout of the hormone from the medium, while T-47D and MDA-MB-231 have had intracellular E2 retention (Fig. 1B). Additionally, MCF-7 cells were able to accumulate E2 above treatment concentrations at lower E2 treatment levels (100 pM), possibly indicating a saturable system of intracellular hormone accumulation (Fig. S1A). E2 accumulation saturated rapidly in MCF-7 cells at around 40 minutes of 100 nM E2, with a somewhat slower saturation at 100 nM E2 (Fig. S1B). Interestingly, MCF-7 cells retained a substantial amount of hormone (>7 nM E2) even after 24 hours of washout (Fig. S1C).

**Figure 1.**
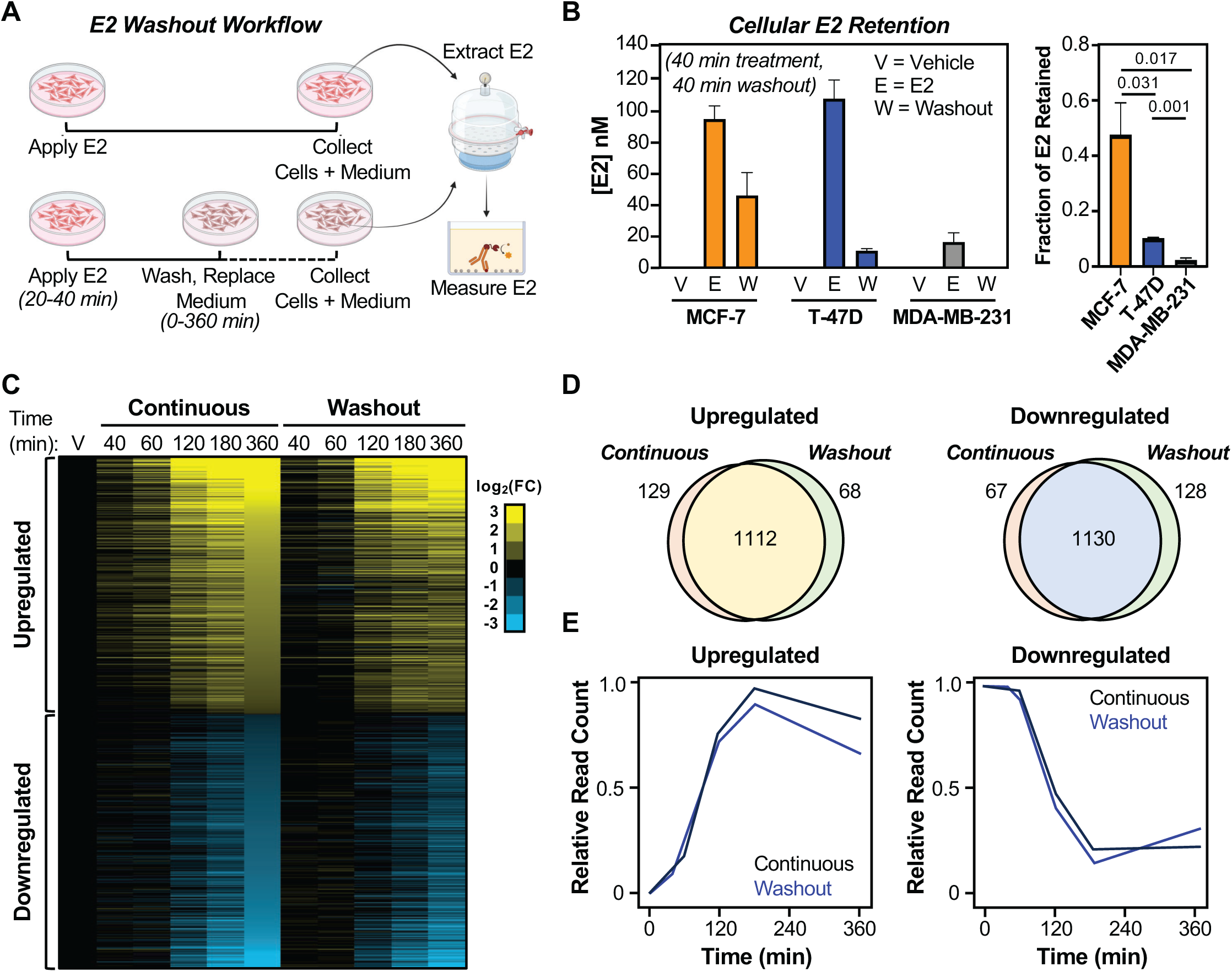
Intracellular E2 retention in MCF-7 cells is associated with sustained gene expression after hormone removal. **(A)** Schematic showing workflow of induction and washout experiments. **(B)** Cellular E2 levels of breast cancer cell lines including MCF-7, T-47D, and MDA-MB-231 as measured by an ELISA-based assay. Initial treatment with E2 was 40 min, followed by 20 min of washout. Fraction of E2 retained in cells were calculated by dividing measured E2 levels in the washout conditions with their respective E2 treated conditions. Significance was determined using multiple Student’s t-tests (non-parametric) with a Šidák-Bonferroni correction. p-values are indicated, n = 3. **(C)** Heatmap showing fold-change gene expression derived from RNA-seq of MCF-7 cells for continuous treatment and washout experiments. Initial treatment with E2 was 40 min, followed by 20 min of washout. Genes are ranked based on fold change at endpoint (360 mins) of continuous treatment conditions. **(D)** Venn diagrams showing the overlap of all differentially expressed genes between the continuous treatment and washout conditions for both upregulated (*left panel*) and downregulated (*right panel*) genes. **(E)** Graph showing relative reads per kilobase mapped reads (RPKM) for continuous treatment and washout conditions across different timepoints for upregulated (*left panel*) and downregulated (*right panel*) genes.

In order to test whether retained E2 is conducive to cellular signaling, we examined changes in gene expression. Remarkably, global gene expression was similar between conditions of continuous E2 treatment and washout conditions at least six hours post-treatment for both upregulated and downregulated genes (Fig. 1C). Furthermore, the total number of genes regulated by E2 in both continuously-treated and washout conditions were similar for each timepoint (Fig. 1D and 1E), suggesting sustained E2 induction hours after removal of hormone from the culture medium. Gene ontology analysis indicated functional similarities between the genes regulated in the continuous treatment and washout groups (Fig. S1D), which is likely due to the fact that the genes sets are very similar.

To further determine how washout of hormone affects E2 the induction of gene expression over time, we categorized the upregulated genes into bins based on the time point of maximum fold change in expression versus vehicle control (Bin 1 = 40 min, Bin 2 = 1 hr, Bin 3 = 2 hr, Bin 4 = 3 hr, and Bin 5 = 6 hr) (Fig. 2A and 2B). In general, number of genes increased in each subsequent bin (Fig. S2A), with the extent of gene expression similar in the washout condition compared to continuous treatment for Bins 1, 4, and 5 (Fig. 2A and 2C). However, gene expression in Bins 2 and 3 was lower in the washout condition as compared to continuous treatment (Fig. 2A and 2C). This suggests that genes with early induction are more sensitive to the removal of hormone.

**Figure 2.**
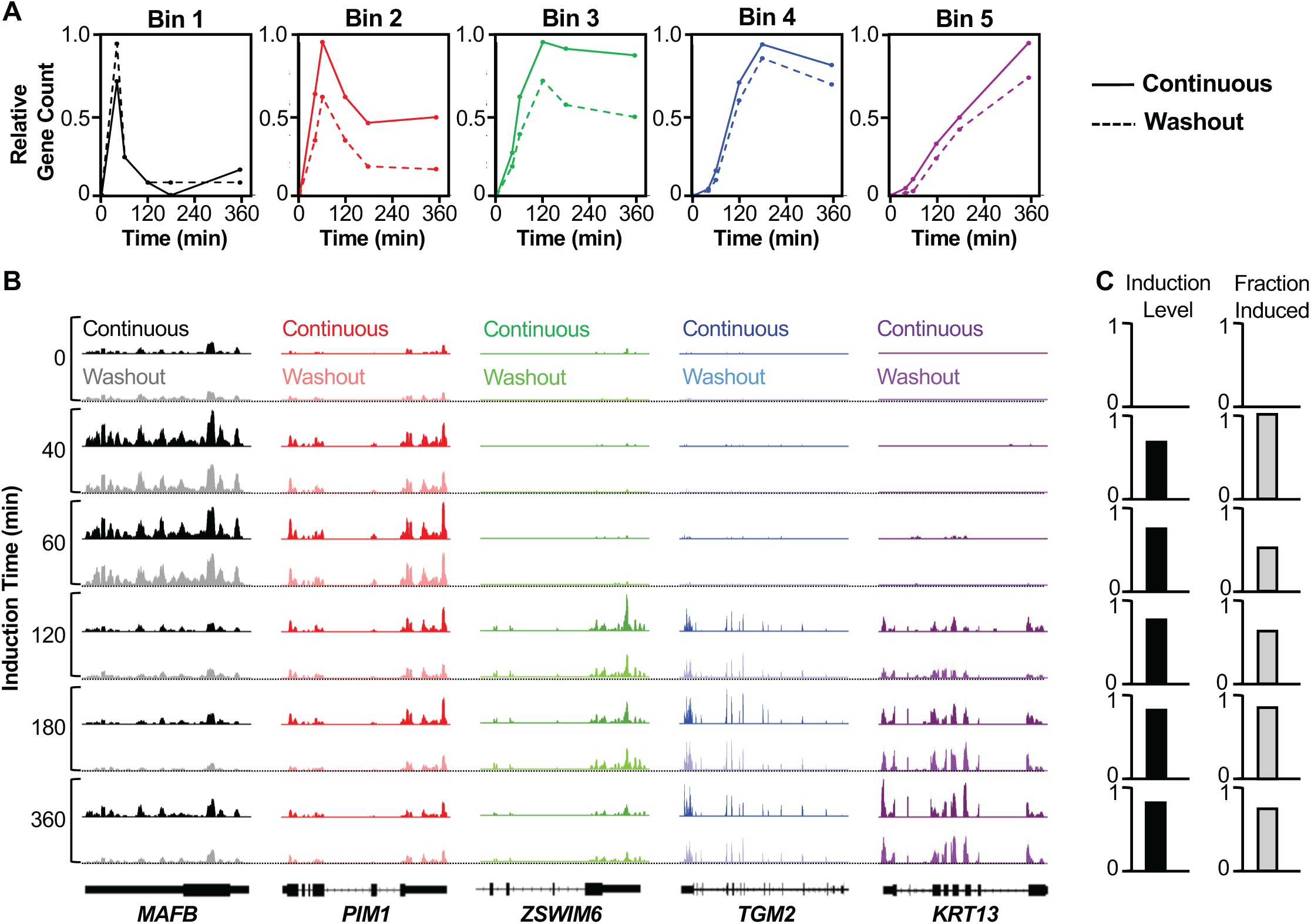
Analysis of steady-state gene expression upon continuous E2 treatments or after washout. **(A)** Bins showing relative number of genes grouped based on time of maximum expression for both continuous and washout groups (Bin 1 = 40 min., Bin 2 = 1 hr., Bin 3 = 2 hr., Bin 4 = 3 hr., and Bin 5 = 6 hr.). Solid lines represent continuous treatment and dotted lines represent washout treatment. **(B)** Browser tracks of representative genes for each bin. From left to right black = Bin 1, red = Bin 2, green = Bin 3, blue = Bin 4, and purple = Bin 5. Darker colors represent continuous treatment and lighter colors represent washout treatment. **(C)** Graphs showing induction levels calculated by relative number of genes in washout group with fold-change induction >0.8 that of continuous treatment groups for each bin or fraction of genes induced calculated by the ratio of genes in the washout to continuous treatment that have fold-change induction >1.5.

In order to assess whether differences between sensitivity to hormone removal were due to a decrease in overall fold-change gene expression or overall change in number of genes, we determined induction levels and the fraction induced across the bins. Induction levels were determined as fraction of genes with induction levels in the washout at least 0.8-times that of continuous treatment. Induction levels of >0.8 were seen in each bin, indicating that the genes are expressed in washout groups to the same extent as the continuous treatment group (Fig. 2C). However, when assessing the relative number of genes in washout to continuous groups in each bin, fewer genes were present in bins 1 and 2 (Fig. 2C). Taken together, the differences between washout and continuous induction at the early timepoints (Bins 1 and 2) were due to fewer number of induced genes in the washout groups rather than differences in the levels of gene expression.

Finally, analysis of motifs at nearest neighboring enhancers based on previously published ChIP-seq data (18) showed changes in the top motif usage across the bins (Fig. S2B and S2C). For example, although EREs were enriched in both conditions across all bins, motifs for the pioneer factors FOXA1 and GATA3 showed differential enrichment in the washout condition (Fig. S2B and S2C), suggesting a link between induction time and sequence motif usage.

### Non-steroidal ER**α** ligands do not affect E2 accumulation or retention

Given that extracellular circulating steroid hormones are sequestered by SHBG (10), we hypothesized that intracellular retention of E2 may also be working through its steroidal properties. Cells treated acutely with fluorescently labeled estradiol (E2-Glow) for 20 minutes accumulated and retained a substantial amount of hormone intracellularly (Fig. 3A). Retained E2-Glow was resistant to washout with hormone-free medium for over 20 min (Fig. 3A). Additionally, unlabeled E2 efficiently competed with intracellular E2-Glow, while the non-steroidal ERα ligands diethylstilbestrol (DES) and 4-hydroxytamoxifen (4-OHT) failed to compete (Fig. 3B). These results indicate that accumulation and retention of E2 are due to its steroidal properties.

**Figure 3.**
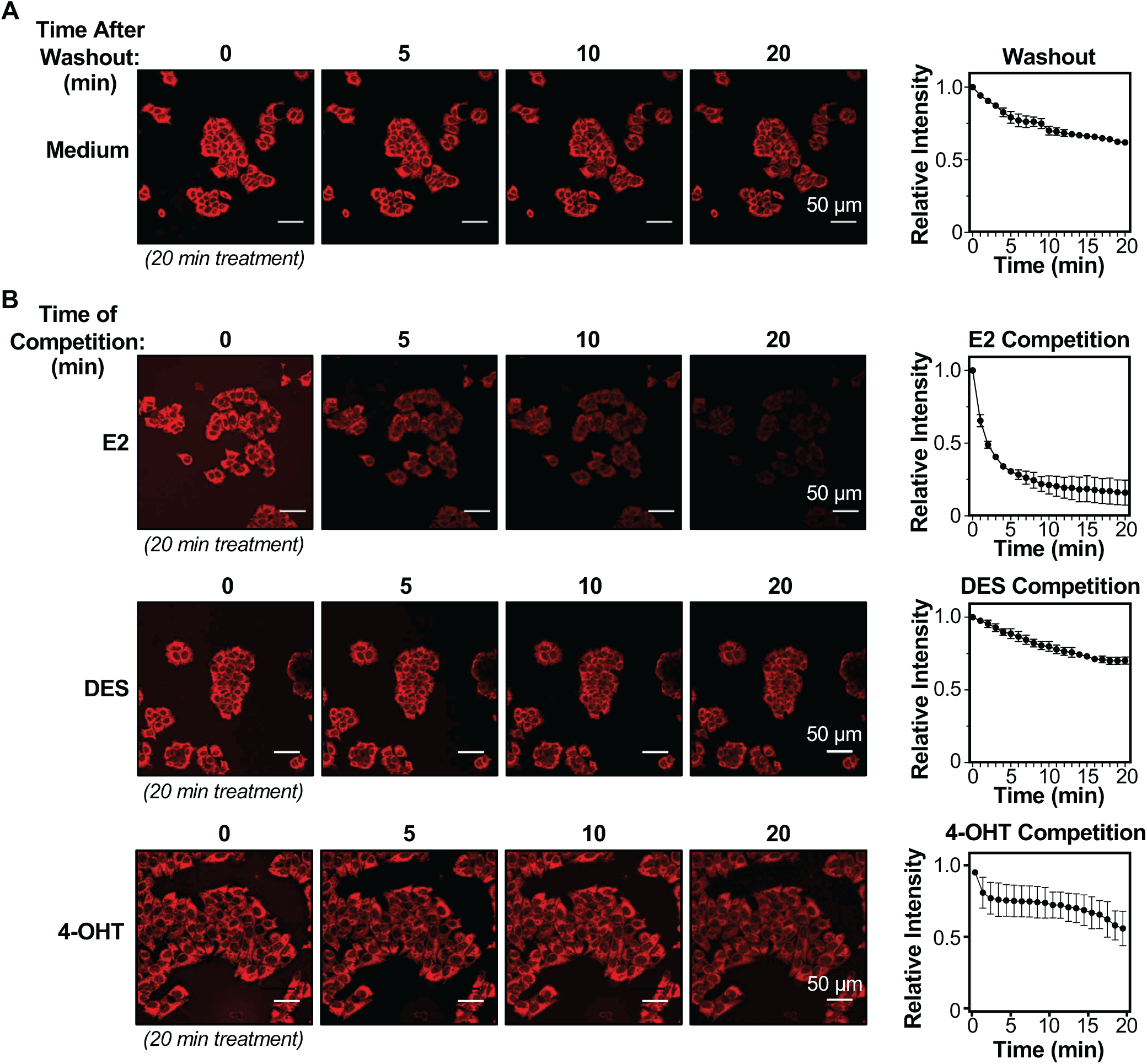
Washout and competition analyses of E2 retention in MCF-7 cells. **(A)** Washout with hormone-free stripped medium after acute treatment with 100 nM E2-Glow reagent for 20 min followed by 20 min of washout. Time t = 0 min indicates start of washout. Quantification of relative intensity of washout experiment images from n = 3 independent biological replicates normalized to value at t = 0 mins (time of washout). The scale is indicated. **(B)** Competition with 100X concentration of unlabeled E2 (*top panel*), 100X DES (*middle panel*), or 4-OHT (*bottom panel*) after acute treatment with 100 nM E2-Glow for 20 min. Time t = 0 min indicates time of addition of competitor. Quantification of competition experiment images from n = 3 independent biological replicates normalized to value at t = 0 mins (time after addition of unlabeled E2, DES, or 4-OHT). The scale is indicated.

### ASTER-B is aberrantly localized into the nucleus of some breast cancer cells

ASTER-B has been recently reported to be involved in E2 homeostasis (16). Promiscuous binding of ASTER-B to cholesterol-derived molecules, including E2, led us to investigate the role of ASTER-B in intracellular E2 retention. ASTER-B protein is expressed at varying levels in a set of ERα-positive and ERα-negative breast cancer cell lines (Fig. 4A). While ASTER-B normally localizes to the cytosol on the endoplasmic reticulum through its transmembrane domain, recent studies have shown that it can aberrantly translocated into the nucleus of certain gastric cancers (19). In fact, sequence analysis of ASTER-B using a publicly available prediction software (NovoPRO) identified a putative nuclear localization signal (NLS) C-terminal to the VASt/ASTdomain spanning amino acids 550-570 (Fig. 4B) (20). Immunofluorescent staining of a panel of breast cancer cell lines shows that MCF-7 cells and ZR-75-1 cells (both ERα-positive) have aberrantly localized nuclear ASTER-B, while T-47D, MDA-MB-231, and MDA-MB-361 have cytosolic ASTER-B (Fig. 4C).

**Figure 4.**
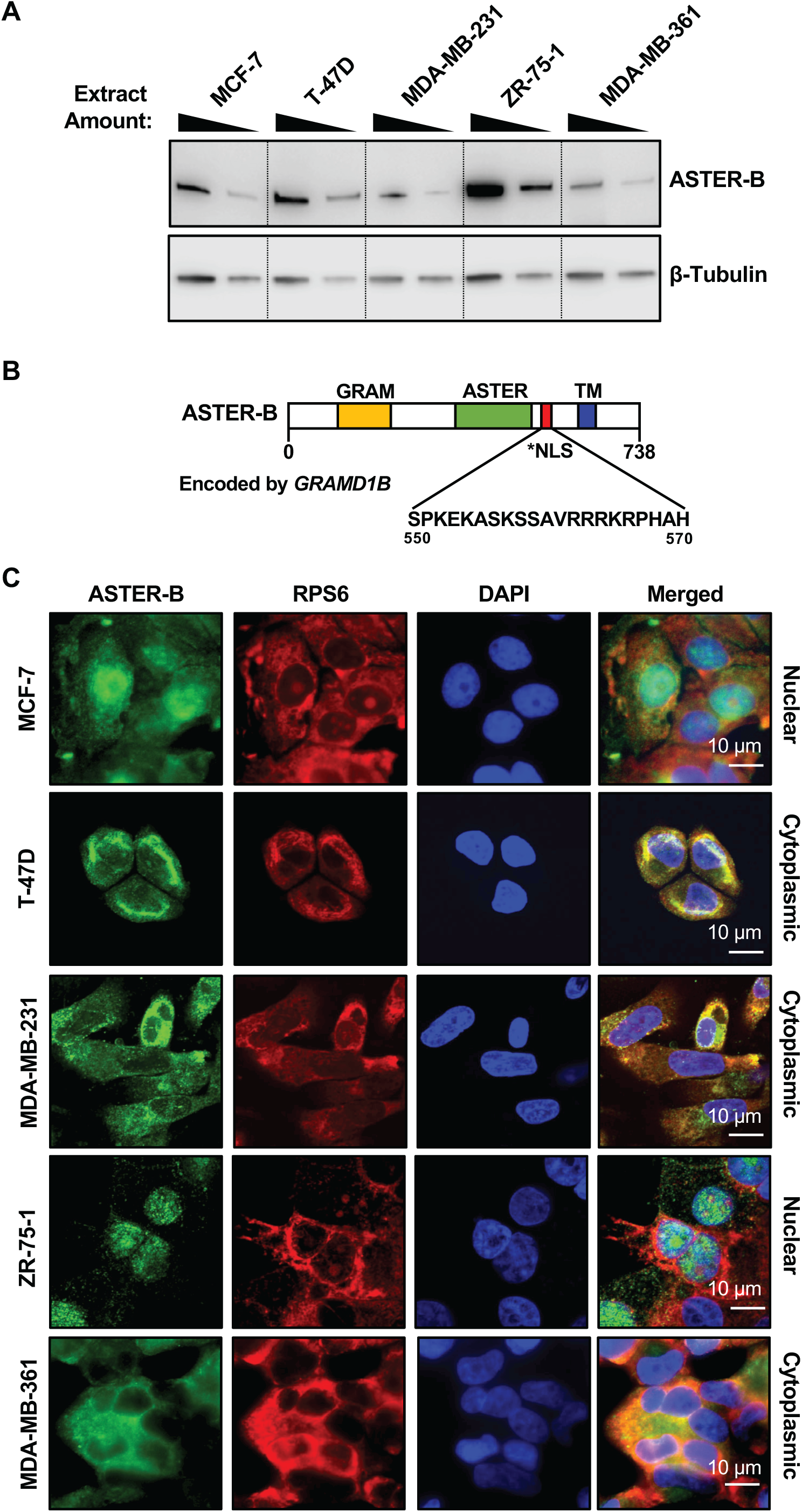
ASTER-B localization in breast cancer cells. **(A)** Western blot showing the expression of ASTER-B across breast cancer cell lines (MCF-7, T-47D, MDA-MB-231, ZR-75-1, and MDA-MB-361) at two dilutions of whole cell extracts. **(B)** Schematic showing the domain structure of ASTER-B protein. Putative nuclear localization signal (NLS) is highlighted in red along with its sequence. **(C)** Immunofluorescence images of breast cancer cell lines stained for ASTER-B (*green*), RPS6 endoplasmic reticulum stain (*red*), and DAPI nuclear stain (*blue*). The scale is indicated.

### Steroid binding and nuclear localization by ASTER-B promote the cellular retention of E2

Previous studies examining the ligand-binding abilities of ASTER-family proteins revealed a G518F mutation located within the hydrophobic cavity in ASTER-B that greatly reduces its ability to bind a variety of cholesterol-derived molecules (21). In order to better understand how steroid binding and nuclear localization by ASTER-B affects intracellular E2 retention, we generated wild-type (WT) and mutant (G518F) constructs with deletion of the transmembrane domain or the addition of a strong nuclear localization signal (NLS) at the N-terminus (Fig. 5A). We also generated a mutant with deletion of the putative endogenous NLS in order to modulate the subcellular localization (Fig. 5A). These proteins were expressed ectopically as FLAG-tagged constructs in HEK-293T cells, and expression and subcellular localization were determined by Western blotting and immunofluorescence analysis, respectively (Fig. 5, B through D). As expected, full length ASTER-B (FL) and ASTER-B with deletion of the putative NLS (ΔTM-ΔNLS) were localized primarily to the cytoplasm/ endoplasmic reticulum, as indicated by co-staining with endoplasmic reticulum marker RPS6 (Fig. 5D). This confirms that ASTER-B (FL) has mainly cytosolic localization and demonstrates that the putative NLS can support nuclear localization of ASTER-B. Expression of ASTER-B with an N-terminal NLS, deletion of the transmembrane domain (ΔTM), or both resulted in mainly nuclear localization (Fig. 5D). This indicates that subcellular localization can be modulated by addition of a localization signal or by removal of the TM domain, which anchors ASTER-B to the endoplasmic reticulum.

**Figure 5.**
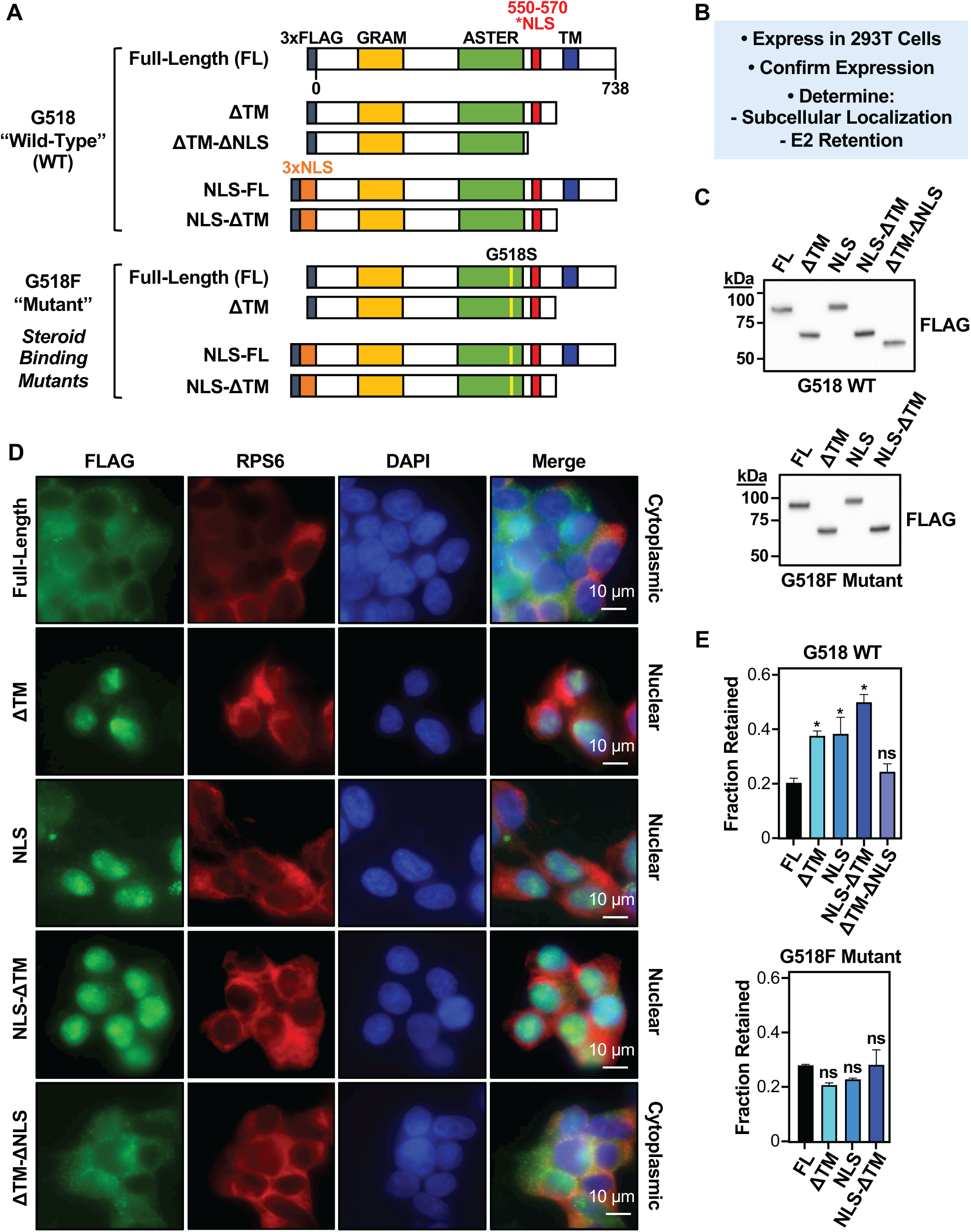
Effects of ectopic expression of ASTER-B mutant proteins on the cellular retention of E2. **(A)** Schematic diagram showing the structure of FLAG-tagged ASTER-B wild-type and G518F steroid binding deficient mutant constructs used in the experiments outlined in **(B)**. **(C)** Western blots showing the size and expression of the ASTER-B G518 wild-type (*top*) and G518F mutant (*bottom*) proteins. **(D)** Immunofluorescent staining showing localization of FLAG-tagged ASTER-B constructs (green), RPS6 endoplasmic reticulum stain (red), and DAPI nuclear stain (blue) in HEK-293T cells. The scale is indicated. **(E)** Cellular E2 levels of HEK-293T cells expressing ASTER-B wild-type (*top*) or mutant (*bottom*) constructs as measured by an ELISA-based assay. The fraction of E2 retained in cells was calculated by dividing the measured levels of E2 in the washout conditions with their respective E2-treated conditions. Significance was determined using multiple Student’s t-tests (non-parametric) with a Šidák-Bonferroni correction Significance values were assigned as follows: * p<0.033, n = 2.

Next, we tested whether ectopic expression of these constructs could augment the cellular retention of E2. HEK-293T cells ectopically expressing the various wild-type and mutant ASTER-B proteins were subjected to acute E2 treatment followed by washout, and intracellular E2 was measured as described above. We observed that expression of mutant ASTER-B proteins that localize to the nucleus increased cellular E2 retention significantly more than the ASTER-B proteins that localize to the cytoplasm (Fig. 5E). Furthermore, none of the G518F mutant constructs enhanced E2 retention regardless of localization, demonstrating the critical role for steroid-binding as a key determinant for E2 retention (Fig. 5E). Taken together, our results indicate that cellular retention of E2 is augmented by both ASTER-B subcellular localization, as well as its steroid-binding ability.

### Nuclear localization of ASTER-B supports E2-dependent gene regulation and cell growth

Given ASTER-B’s ability to bind and accumulate E2 in cells, we explored whether its expression and nuclear localization may be associated with E2-dependent gene regulation. We knocked down *GRAMD1B* mRNA, which encodes ASTER-B, using siRNAs, which caused a depletion in the levels of ASTER-B protein (Fig. 6A and 6B). Knockdown of *GRAMD1B* mRNA in MCF-7 cells dramatically decreased intracellular E2 accumulation after washout, but had little effect with continuous E2 treatment, without changes in E2 levels in the medium (Fig. 6C and 6D). Knockdown of *GRAMD1B* mRNA in MCF-7 cells also reduced the retention of E2-Glow (Fig. S3A). In parallel, we also observed that the levels of ERα in MCF-7 cells also affect the retention of E2; depletion of ERα reduced E2 retention (Fig. S3B and S3C), whereas ectopic expression of ERα enhanced E2 retention (Fig. S3D and S3E). These results suggest a potential contribution of ERα to E2 retention, at least in cells with very high levels of ERα. Finally, as an addition control experiment, we observed no effect of E2 treatment on the levels of ASTER-B protein, but did observe a typical E2-dependent reduction ERα (Fig. S3F).

**Figure 6.**
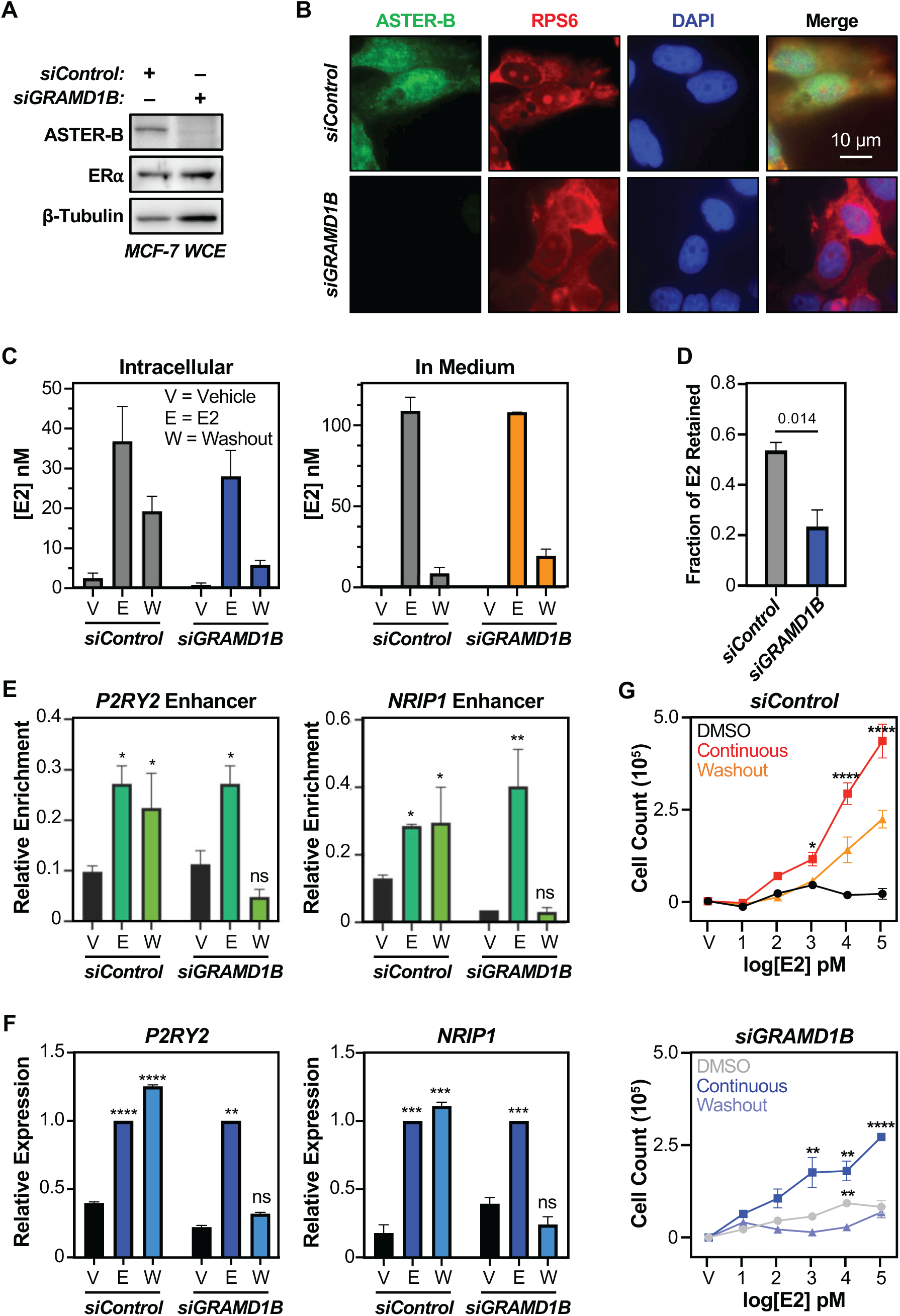
ASTER-B nuclear localization is associated with cellular retention of E2, regulation of gene expression, and growth in MCF-7 cells. **(A and B)** Western blot (A) and immunofluorescence images (B) showing knockdown of ASTER-B in MCF-7 cells. The scale in panel B is indicated. **(C and D)** Measurement of intracellular (*left panel*) and culture medium (*right panel*) E2 levels by ELISA-based quantification. In panel D, significance was determined using multiple Student’s t-tests (non-parametric) with a Šidák-Bonferroni correction. The p-value is indicated, n = 3. **(E and F)** Chromatin occupancy of ERα at enhancer regions and its corresponding gene expression at E2 regulated genes (*P2RY2* and *NRIP1*) measured by ERα ChIP-qPCR and RT-qPCR respectively. For the ChIP-qPCR assays, initial treatment with E2 was 40 min, followed by 20 min of washout. For the RT-qPCR assays, initial treatment with E2 was 40 min, followed by 1 hour of washout. For statistical analysis, comparisons were made against vehicle control for their own treatment groups. Multiple Student’s unpaired t-tests (non-parametric) with a Šidák-Bonferroni correction used to determine significance. Significance values were assigned as follows: * p<0.033, ** p<0.002, *** p<0.0001, **** p<0.00001. n = 3. **(G)** Growth assay of MCF-7 cells treated continuously with 100 nM E2, or treated acutely for 40 mins on day 1 (washout), or treated with vehicle control. The cells were also treated with control siRNA (*top panel*) or si*GRAMD1B* (*bottom panel*). The cell counts were made on day 5 of the assay. Multiple Student’s unpaired t-tests (non-parametric) with a Šidák-Bonferroni correction used to determine significance. Significance values were assigned as follows: * p<0.033, ** p<0.002, *** p<0.0001, **** p<0.00001. n = 3.

We next explored the effects of E2 retention on molecular features of E2 signaling. We observed the maintenance of ERα binding at the enhancer regions of E2-responsive genes (*P2RY2*, *NRIP1*, *TFF1*, and *MBOAT*) were maintained even after the washout of E2 from the culture medium (Fig. 6E, Fig. S4A). The maintenance of ERα binding upon E2 washout, however, was lost when *GRAMD1B* mRNA was knocked down (Fig. 6E, Fig. S4A). Furthermore, the E2-dependent expression the cognate genes reflected the binding of ERα at the enhancers (i.e., loss of E2-dependent expression when *GRAMD1B* mRNA was knocked down) (Fig. 6F, Fig. S4B), indicating that cellular E2 accumulation is necessary for both sustained chromatin binding and gene expression. In cell growth assays with MCF-7 cells, we observed that E2 had a robust mitogenic effect, with significant proliferative effects observed upon either continuous E2 treatment or washout (Fig. 6G, *top*). These effects were inhibited, however, with knockdown of *GRAMD1B* mRNA, with reduced E2-dependent growth observed with continuous E2 treatment and a loss of reduced E2-dependent with washout (Fig. 6G, *bottom*). Taken together, these results demonstrate that ASTER-B expressed from *GRAMD1B* mRNA supports cellular accumulation and retention of E2, which is necessary to sustain gene regulatory programs upon E2 washout that promote a mitogenic response.

### ASTER-B is aberrantly localized to the nucleus in some breast cancers

To determine the broader biological relevance of our observations, we investigated ASTER-B in breast cancer patient samples. We analyzed RNA-seq data from The Cancer Genome Atlas (TCGA) (22) and found that *GRAMD1B* mRNA expression is not significantly associated with prognosis (overall survival) in all breast cancers or the luminal A or luminal B molecular subtypes (Fig. 7A, Fig. S5A). These results fit with our observation that nuclear localization is the key determining factor for ASTER-B-mediated retention and accumulation of E2.

**Figure 7.**
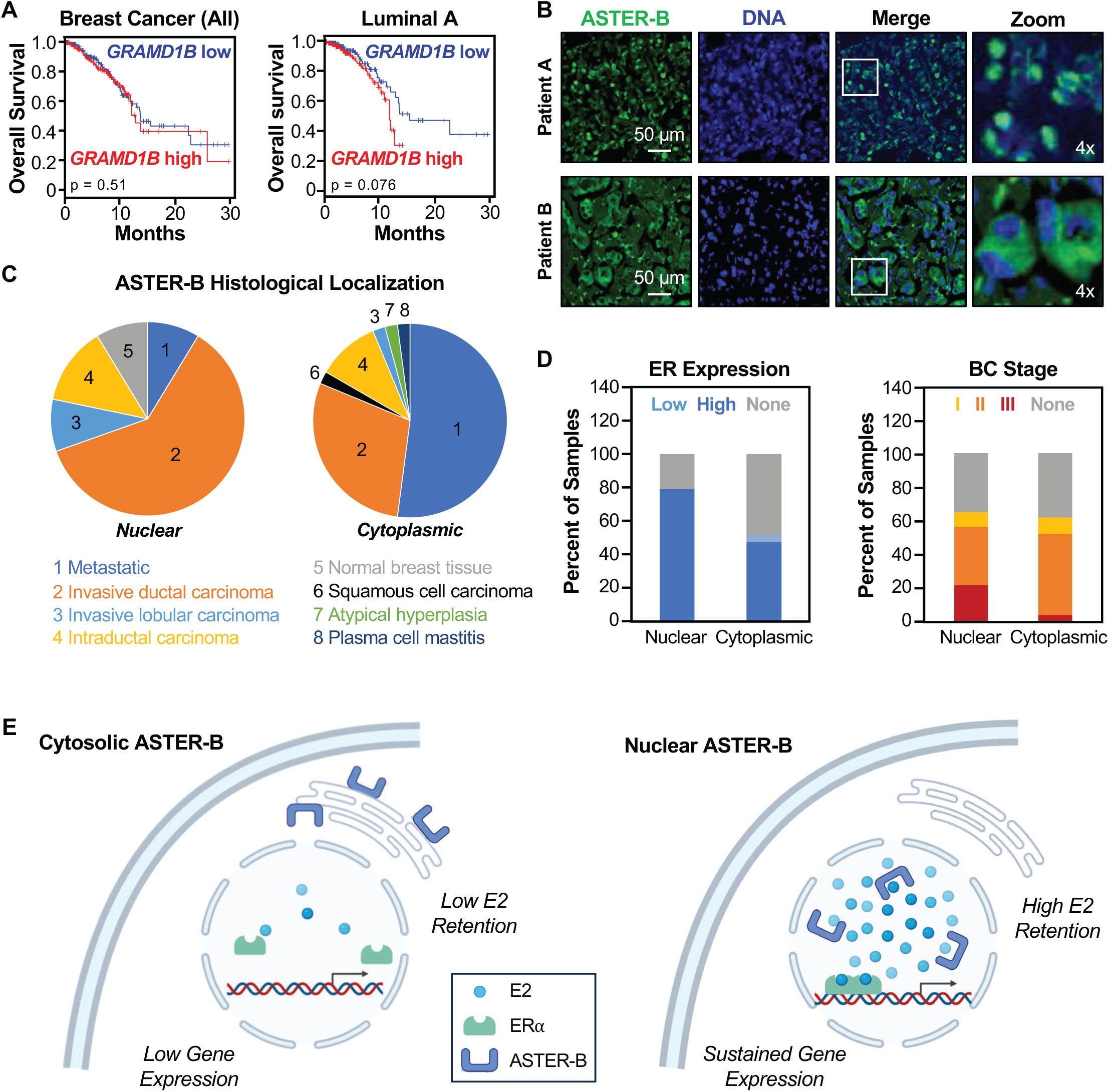
Nuclear ASTER-B localization is associated with decreased survival. **(A)** Kaplan-Meier survival plots of all TCGA breast cancer patients (n = 1064; *left panel*) and specifically luminal A subtype breast cancer patients (n = 415; *right panel*) with high (*red line*) or low (*blue line*) *GRAMD1B* mRNA expression level over a 5-year period. P-values were calculated using the Log-rank test. **(B)** Representative images of either nuclear (*top panel*) or cytoplasmic (*bottom panel*) ASTER-B (*green*) co-stained with DAPI (*blue*). **(C)** Pie charts showing the histological subtypes of patients with either nuclear localized (*left panel*) or cytoplasmic (*right panel*) ASTER-B as assessed by immunostaining of a commercial breast cancer tissue microarray. **(D)** Graphs showing ERα expression status (*left panel*) and breast cancer stages (*right panel*) of nuclear or cytoplasmic ASTER-B in a breast cancer tissue microarray as determined by immunohistochemistry (IHC) staining and histology respectively. **(E)** Schematic showing a working model of ASTER-B mediated E2 cellular retention. The schematics were generated using BioRender (BioRender.com). Details are provided in the text.

In this regard, we performed immunostaining for ASTER-B using a breast cancer tissue microarray, which identified samples with either nuclear or cytoplasmic ASTER-B localization (Fig. 7B; Fig. S5B). We observed nuclear localized ASTER-B in 24 out of 192 samples (13%), while 50 out of 192 samples (26%) had cytoplasmic localization and 118 out of 192 samples (61%) had no significant expression (Fig. 7C). Of the patient samples with nuclear ASTER-B localization, a majority (61%) were classified as invasive ductal carcinoma (Fig. 7C). In contrast, the patient samples with cytoplasmic ASTER-B localization were mostly of metastatic (52%). Additionally, a higher proportion of samples with nuclear ASTER-B had high ERα expression, as well as a higher proportion of Stage III breast cancers, as compared to samples with cytosolic ASTER-B (Fig. 7D). Collectively, our results indicate that ASTER-B is aberrantly localized to the nucleus in both cultured cells and breast cancer patient samples. In patients, nuclear localization of ASTER-B is positively associated with ERα expression and breast cancer progression.

## Discussion

In this study, we explored the association between a cholesterol binding protein, ASTER-B, and intracellular retention of E2. We showed that MCF-7 cells accumulate and retain E2 for a prolonged duration even after hormone removal (Fig. 1A and 1B; Fig. S1A-S1C). Retained E2 is associated with sustained cell proliferation (Fig. 6G), ERα enhancer activity (Fig. 6E; Fig. S4A), and E2-dependent gene expression(Fig. 6F; Fig. S4B). We observed aberrant nuclear localization of ASTER-B in MCF-7 cells and ZR-75-1 cells, which was associated with E2 retention, but not in some other commonly used breast cancer cell lines (Fig. 4). In our experiments, we determined that both the mislocalization of ASTER-B, as well as ASTER-B’s steroid-binding ability, were required for the retention and accumulation of E2 (Fig. 5). Aberrant nuclear localization of ASTER-B was also observed in tumor samples from a subset of breast cancer patients (Fig. 7B-7D). Moreover, depletion of ASTER-B by knocking down of *GRAMD1B* mRNA, which encodes ASTER-B, led to a significant decrease in cellular E2 retention, gene regulation (i.e., ERα binding at enhancers and E2-dependent gene expression), and cell proliferation (Fig. 6). Collectively, our results indicate that mislocalization of ASTER-B to the nucleus can provide an alternate means of enhancing the biological effects of E2 in breast cancers.

### Mislocalization of ASTER-B leads to intracellular E2 retention

Little is known about how estrogen may be retained within the cell or how this may affect hormone-dependent gene regulation. We showed that acute treatment can support both cell proliferation as well as gene expression. Based on our genomic and cell-based assays, we find that ASTER-B mislocalization leads to increased intracellular E2 retention. This in turn supports ERα binding, gene expression, and cell proliferation for a prolonged duration after removal of E2 from the culture medium. While the mechanism of nuclear translocation of ASTER-B is not fully understood, it occurs in a variety of different pathologies, including gastrointestinal cancers and breast cancer (19). Initial analysis of COSMIC database shows mutations within the transmembrane domain of ASTER-B that may allow for nuclear translocation through its cryptic NLS. Our results with mutant ASTER-B proteins targeting the NLS suggest that the cryptic NLS does, indeed, play a key role in the nuclear localization. Comparison between different breast cancer cell lines indicates that the nuclear localization of ASTER-B, and not necessarily its expression, that contributes to E2 retention.

### ASTER-B supports E2-mediated gene regulation

We have demonstrated that hours after treatment, E2-regulated genes are expressed at the same fold-change intensity as continuous treatment. These effects are diminished, along with ERα binding at its enhancers, with the knockdown of ASTER-B and the reduction of intracellular E2 accumulation. These results suggest that sustained induction by E2 is contingent on intracellularly available estrogen. Additionally, a recent study has shown that ASTER-B is involved in supporting E2 synthesis in the ovaries (16). Given its dual role in both estrogen production and intracellular retention, ASTER-B may be a potential therapeutic target that addresses multiple aspects of estrogen bioavailability.

### Clinical implications of ASTER-B mislocalization

Analysis of tissue samples from ER+ breast cancer patients showed that a subset of patients with nuclear ASTER-B localization. These patients had a modest increase of both ERα expression, as well as poorer prognosis. Combined with the cell-based assays performed in this study, these results suggest that elevated E2 in the tumors of these patients may contribute to the outcomes. It is plausible that this may provide a growth advantage for ER+ breast cancer even in environments of fluctuating or low estrogens. In fact, the highest instances of ER+ breast cancer are in postmenopausal women, where circulating E2 levels are greatly diminished (23). An increased ability to not only retain, but accumulate, estrogens may be highly relevant to tumor survival and disease progression. This may be especially important in the tumor microenvironment, with multiple cells that may have different abilities to retain and exchange E2. In this regard, *GRAMD1B* mRNA is expressed across a wide range of cell types, some of which may represent a component of the tumor microenvironment for breast cancers (Fig. S5C).

Additionally, we found that cells with nuclear ASTER-B are resistant to the displacement of E2 by non-steroidal ERα ligands, including 4-OHT, which is used in the treatment of ER+ breast cancer. In this regard, treatments that act as competitive inhibitors of E2 binding to ERα may be less effective in cells with nuclear-localized ASTER-B. Additionally, ASTER-B-mediated E2 accumulation may support growth of ER+ breast cancers despite suppression of circulating hormone by aromatase inhibitors. This, ASTER-B nuclear localization status in breast cancer could be an important consideration for therapeutic efficacy of ERα-targeted therapeutics (Fig. 7E).

## Materials and Methods

### Antibodies

The custom rabbit polyclonal antiserum against ERα was generated, as previously described (24). The other antibodies used were as follows: rabbit polyclonal anti-GRAMD1B (ASTER-B) (Proteintech, 24905-1-AP; RRID:AB_2879791), mouse monoclonal anti-RPS6 (Santa Cruz, sc-74459; RRID:AB_1129205), rabbit polyclonal anti-DYKDDDDK (Invitrogen PA1-984B), and rabbit polyclonal anti-β-tubulin (Abcam, ab6046; RRID:AB_2210370).

### Cell culture

MCF-7 cells were kindly provided by Dr. Benita Katzenellenbogen (University of Illinois at Urbana-Champaign, IL) and were maintained in minimal essential medium (MEM; Sigma M1018) supplemented with 5% calf serum (Sigma, C8056), 100 units/mL penicillin-streptomycin (Gibco, 15140122), and 25 μg/mL gentamicin (Gibco, 1571004). All other cell lines were obtained from the American Type Culture Collection (ATCC). MDA-MB-231 and HEK-293T cells were maintained in Dulbecco’s modified Eagle’s medium-low glucose (DMEM; Sigma, D6046) supplemented with 10% fetal bovine serum (Sigma, F0926), 2 mM GlutaMAX (Gibco, 35050061), 100 units/mL penicillin-streptomycin (Gibco, 15140122), and 25 μg/mL gentamicin (Gibco, 1571004). T-47D and ZR-75-1 cells were maintained in Roswell Park Memorial Institute medium (RPMI; ThermoFisher, 11875119) supplemented with 10% fetal bovine serum (Sigma, F0926), 2 mM GlutaMAX (Gibco, 35050061), 100 units/mL penicillin-streptomycin (Gibco, 15140122), and 25 μg/mL gentamicin (Gibco, 1571004). MDA-MB-361 cells were maintained in Alpha-MEM medium (Sigma, M8042) supplemented with 0.1 M HEPES, 10% fetal bovine serum (Sigma, F4135), 1% non-essential amino acids, 2 mM L-glutamine, 1% sodium pyruvate, 1 µg/mL insulin, 1 ng/mL hydrocortisone, 12.5 ng/mL epidermal growth factor, and 1% penicillin/streptomycin. All cell lines were authenticated for cell type identity using the GenePrint 24 system (Promega, B1870), and confirmed as *Mycoplasma*-free every six months using the Universal Mycoplasma Detection Kit (ATCC, 30-1012K). Fresh cell stocks were regularly replenished from original stocks every few months (no more than 10 passages).

### Estradiol treatment and washout experiments

Seventy-two hours prior to E2 treatment, cells were cultured in hormone-free stripped medium consisting of Eagle’s phenol red-free minimal essential medium (Sigma, M3024) supplemented with 5% charcoal-dextran treated calf serum (CDCS, Sigma, C8056), 2 mM GlutaMAX, 100 units/mL penicillin-streptomycin, and 25 μg/mL gentamicin. All subsequent experiments involving E2 induction were carried out in stripped medium. For continuous induction experiments, cells were treated with DMSO (vehicle) or 100 nM 17β-estradiol (E2) (Sigma, E8875) for the indicated times. For washout experiments, cells were induced for 40 min. with vehicle or 100 nM E2, washed once with stripped medium, then incubated in stripped medium for the remainder of the indicated time.

### siRNA knockdown

For siRNA-mediated knockdown, siRNA against *GRAMD1B* (SASI_Hs02_0035373), *ESR1* (SASI_Hs02_00078592), or scramble siRNA (Sigma, SIC001) were purchased from Sigma and transfected into MCF-7 cells at a final concentration of 30 nM by electroporation using the SE Cell Line 4D-Nucleofector X kit (Lonza; EEID:SCR_023155). The cells were used for various assays 48 hours after siRNA transfection.

### Ectopic expression of ER**α** and ASTER-B

For experiments involving ectopic expression of ERα or ASTER-B, HEK-293T cells were transfected with mammalian expression vectors for ERα (pCMV-ERα) (25) or the ASTER-B expression constructs described below using Lipofectamine 3000 (ThermoFisher, L3000015) following the manufacturer’s protocol. Cells were transfected 48 hours prior to experimentation.

### Generation and expression of modified and mutant ASTER-B proteins

Mammalian expression vectors for wild-type FLAG-tagged ASTER-B (encoded by the *GRAMD1B* cDNA), truncation/deletion constructs, and corresponding G518F mutants were generated for ectopic expression. A full length *GRAMD1B* cDNA in the pcDNA3.1-C-(k)DYK vector was obtained from Genscript (OHu11754) and cloned into pcDNA3 (Addgene, 58792) with an N-terminal 3X FLAG-tag. Truncation/deletion and NLS subclones were generated by PCR amplification from the full-length FLAG-tagged construct, followed by Gibson assembly using the HiFi DNA Assembly Master Mix (NEB, E2621S) in pcDNA3 vectors that were linearized by digestion with *XhoI* (NEB, R0146S) and *HindIII* (NEB, R0104S). The G518F mutation was introduced into the pcDNA3-FLAG-*GRAMD1B* construct using the Q5 Site-Directed Mutagenesis Kit (NEB, E0554S) following the manufacturer’s protocol. NLS and truncation/deletion mutant subclones were generated for G518F mutant constructs using the same amplicon primers and assembly strategy as the wild-type constructs detailed above.

### Cloning primers for the ASTER-B expression constructs

The following primers were used to generate FLAG-tagged WT and G518F constructs.

- Forward (3X FLAG): 5’-CGACTCACTATAGGGAGACCCGCCACCATGGACTACAAGGACCACGACGGTGACTACAAGGACCACGACATCGACTACAAGGACGACGACGACAAGAAAGGATTCAAGCTCTCCTGCACTG-3’
- Forward (3X FLAG-3X NLS): 5’-CTCACTATAGGGAGACCCGCCACCATGGACTACAAGGACCACGACGGTGACTACAAGGACCACGACATCGACTACAAGGACGACGACGACAAGCCAAAAAAGAAGAGAAAGGTAGACCCAAAGAAAAAACGAAAAGTAGATCCGAAAAAGAAGAGGAAGGTGAAAGGATTCAAGCTCTCC-3’
- Reverse (Full length): 5’-CTAGCATTTAGGTGACACTATAGAATATCAATGATAGCGATTCCTCTTTTCTTCACTTTC-3’
- Reverse (ΔTM): 5’-CTAGCATTTAGGTGACACTATAGAATATCACTCCTGGCTCGCCTTGTACATGGTCTG-3’
- Reverse (ΔTM-ΔNLS): 5’-GCATTTAGGTGACACTATAGAATATCAGAGCTTGTCCACGCTGAAGTTG-3’

### Primers for introducing the G518F point mutant

The following primers were used to introduce the G518F point mutant using the Q5 Site-Directed Mutagenesis Kit.

- Forward: 5’-CTTCTGGAGTAGCCTGGAGGACTAC-3’
- Reverse: 5’-TTCTTCTCGATGAACGTTTTC-3’

### Measurement of E2 levels by ELISA

Treated cells and culture medium used for induction experiments were collected both before and after experimentation. In short, cells were harvested by washing once in ice cold 1X PBS and collected by scraping. Cells were then pelleted by centrifugation and packed cell volumes (PCV) were estimated. E2 was then extracted from samples by applying 5X volume (PCV or volume of medium) of pure methanol. Samples were vortexed for more than 30 seconds, then incubated at room temperature rotating end-over-end for 15 mins. Insoluble material was pelleted by centrifugation. The supernatant fraction was transferred to a new tube and dried in a vacuum desiccator. The resulting material was resuspended in appropriate volume of ELISA Buffer (provided by the Cayman Chemicals Estradiol ELISA kit). E2 concentrations were measured using the Estradiol ELISA kit (Cayman Chemicals, 501890) as described by the manufacturer’s protocol.

### Experiments with fluorescent E2 (E2-Glow)

Stripped MCF-7 cells were seeded onto poly-lysine, γ-irradiated dishes (MatTek, P35GC-1/5-14-C) one day prior to imaging. Cells were treated with 100 nM Estradiol Glow (Jena Biosciences, PR-958S) for 20 mins at 37°C and images were acquired using an inverted Zeiss LSM 780 confocal microscope at 37°C and 5% CO_2_. For washout experiments, preincubated cells were washed once with stripped medium, then incubated in stripped medium for the indicated time. For competition assays, unlabeled E2 (10 μM), diethylstilbestrol (DES, 10 μM; Cayman Chemicals, 10006876), 4-hydroxytamoxifen (4-OHT, 10 μM; Tocris, Cat. No. 0999) was added directly to the culture medium in vehicle for the indicated time. Images were acquired using the Hamamatsu ORCA-Fusion C14440 digital camera and a CSU-W1 Yokogawa Spinning Disk Field Scanning Confocal System equipped with a Super Resolution by Optical Pixel Reassignment (SoRa) module. Images were acquired using a 20x Plan Apo Lambda objective (NA, 0.75) and a Z-stack step size of 0.2 μm. Exposure time and laser intensity were kept consistent between samples. Laser lines 488 nm and 561 nm were used.

### Identifying a putative NLS

The ASTER-B amino acid sequence was analyzed using a publicly available software from NovoPRO (https://www.novoprolabs.com/tools/nls-signal-prediction) (20). ASTER-B protein sequence was analyzed using the two-state model preset, which uses a hidden Markov model to predict putative NLS. The top NLS signal with >90% probability was reported here.

### Immunofluorescent staining

Breast cancer cells were seeded onto 4-well chambered slides (ThermoFisher, 154917) one day prior to experimentation. The cells were washed once with PBS, fixed with 4% paraformaldehyde for 15 mins at room temperature, and washed three times with 1X PBS. The cells were permeabilized with 0.05% Triton X-100 in 1X PBS for 15 mins at room temperature, washed three times with PBS, and incubated for 1 hour at room temperature in Blocking Solution (1X PBS containing 1% BSA, 10% FBS, 0.3M glycine and 0.1% Tween-20). Fixed cells were incubated with a mixture of the primary antibodies in 1X PBS overnight at 4°C, followed by three washes with PBS. The cells were then incubated with a mixture of Alexa Fluor 594 donkey anti-mouse IgG (ThermoFisher, A-21203; RRID:AB_2535789) and Alexa Fluor 488 goat anti-rabbit IgG (ThermoFisher, A-11008; RRID:AB_143165) each at a 1:500 dilution in 1X PBS for 1 hour at room temperature. After incubation, the cells were washed three times with 1X PBS. Finally, coverslips were placed on cells coated with VectaShield Antifade Mounting Medium with DAPI (Vector Laboratories, H-1200; RRID:AB2336790).

### Preparation of whole cell lysates

Cells were cultured and treated as described above before the preparation of cell extracts. At the conclusion of the treatments, the cells were washed twice with ice-cold 1X PBS and resuspended in lysis buffer (20 mM Tris-HCl pH 7.5, 150 mM NaCl, 1 mM EDTA, 1 mM EGTA, 1% NP-40, 1% sodium deoxycholate, 0.1% SDS) supplemented with 1 mM DTT and 1X complete protease inhibitor cocktail (Roche, 11697498001). The cells were incubated on ice in lysis buffer for 30 mins before vortexing followed by centrifugation at 20,000 g for 15 mins at 4°C in a microcentrifuge to remove the cell debris. Soluble material in the supernatant was collected for further processing.

### Cell growth assays

Cells were maintained in stripped medium for 72 hours prior to growth assay. Cells were plated at a density of 50,000 cells per well in a 6-well dish. Cells were treated either continuously or acutely with washout as indicated above with 100 nM E2. At each 1-day timepoint, cells in a single well were trypsinized and live cell count obtained by staining with 0.4% trypan blue (Sigma, T8154) and counted using the BioRad TC20 automated cell counter. Culture medium was replenished every 2 days with 100 nM E2 for continuous treatment, or untreated stripped medium for washout conditions. Cell counts were determined on day 5.

### Immunoblotting

The protein concentrations of the cell lysates were determined using bicinchoninic acid (BCA) assay (Pierce, 23228). Volumes of lysates were adjusted to the same protein concentrations with lysis buffer and heated to 95°C for 5 mins after addition of appropriate amount of 4X SDS-PAGE Loading Solution (250 mM Tris, pH 6.8, 40% glycerol, 0.04% Bromophenol Blue, 4% SDS). Samples were run on polyacrylamide-SDS gels and transferred to nitrocellulose membranes. After blocking with 5% non-fat milk in TBS-T (50 mM Tris pH7.5, 150 mM NaCl, 0.1% Tween-20), the membranes were incubated with the primary antibodies described above in TBS-T with 0.02% sodium azide, followed by goat anti-rabbit HRP-conjugated secondary antibody at 1:5000 dilution (ThermoFisher, 31463; RRID:AB_228333). Immunoblot signals were detected using an ECL detection reagent (Thermo Fisher Scientific, 34577).

### Reverse transcription quantitative PCR (RT-qPCR)

Breast cancer cells were subjected to siRNA-mediated knockdown and E2 treatment as described above. For the RT-qPCR assays, initial treatment with E2 was 40 min, followed by 1 hour of washout. Total RNA was isolated using the Qiagen RNeasy Plus Mini kit (Qiagen, 74136) according to the manufacturer’s protocol. Total RNA was reverse transcribed using oligo(dT) primers and MMLV reverse transcriptase (Promega, PR-M1705) to generate cDNA. The cDNA samples were subjected to RT-qPCR using gene-specific primers listed below. Target gene expression was normalized to the expression of GAPDH mRNA.

### RT-qPCR primers

The following primer sets were used for RT-qPCR.

- *GAPDH* forward: 5’-CCACTCCTCCACCTTTGAC-3’
- *GAPDH* reverse: 5’-ACCCTGTTGCTGTAGCCA-3’
- *GRAMD1B* forward: 5’-GAGGAGAATGGAAACCAGAGCC-3’
- *GRAMD1B* reverse: 5’-GGTGAGGACTTCGGCATCTATC-3’
- *P2RY2* forward: 5’-CGAGGACTTCAAGTACGTGCTG-3’
- *P2RY2* reverse: 5’-GTGGACGCATTCCAGGTCTTGA-3’
- *NRIP1* forward: 5’-GACCTGGCCCAGATAGATCA-3’
- *NRIP1* reverse: 5’-TATAATGTAGGGGCGCAACC-3’
- *TFF1* forward: 5’-GTTGGGAGCTAGGATGGTCA-3’
- *TFF1* reverse: 5’-AGTGAGTGGCGGATTTGAAC-3’
- *MBOAT1* forward: 5’-GGTTTCCACAGCTTGCCAGAAC-3’
- *MBOAT1* reverse: 5’-ACCAGTCATCCACAAGGCAGGT-3’

### Chromatin immunoprecipitation quantitative PCR (ChIP-qPCR)

Chromatin immunoprecipitation (ChIP) was performed as previously described with some modifications (26). Cells were grown to ~80% confluence and treated as described above. For the ChIP-qPCR assays, initial treatment with E2 was 40 min, followed by 20 minutes of washout. After treatment, the cells were crosslinked with 1% formaldehyde (ThermoScientific, 28-906) in 1X PBS at 37°C for 10 mins. The reaction was then quenched with glycine at a final concentration of 125 mM for 5 mins at 4°C. Cells were washed generously with ice-cold 1X PBS and collected by scraping with 1 mL ice-cold 1X PBS. The cells were pelleted by centrifugation and lysed by pipetting in Farnham Lysis Buffer (5 mM PIPES pH 8, 85 mM KCl, 0.5% NP-40, 1 mM DTT, 1X complete protease inhibitor cocktail). Nuclei were collected by brief centrifugation and resuspended in SDS Lysis Buffer (50 mM Tris-HCl pH 7.9, 1% SDS, 10 mM EDTA, 1 mM DTT, 1X complete protease inhibitor cocktail) by pipetting and incubating on ice for 10 mins. Chromatin was sheared to ~200–500 bp DNA fragments by sonication using a Bioruptor sonicator (Diagenode) for 8 cycles of 30 seconds on and 30 seconds off. Fragment size was verified by agarose gel electrophoresis before quantification of protein concentrations using BCA protein assay kit (Pierce, 23225). Samples were diluted 10-fold using dilution buffer (20 mM Tris-HCl pH 7.9, 0.5% Triton X-100, 2 mM EDTA, 150 mM NaCl) with freshly added 1mM DTT and 1X protease inhibitor cocktail. Soluble chromatin (500 μg per reaction) was precleared with Protein A Dynabeads (Invitrogen, 10001D). Protein A Dynabeads were incubated with 10 μL of polyclonal antiserum for ERα or 5 μg of rabbit IgG control (ThermoFisher, 10500C; RRID:AB_2532981) for 2 hours, washed with dilution buffer, and applied separately to precleared chromatin samples.

The immune complexes from the ChIP were washed once with each of the following in order: (1) low salt [20 mM Tris-HCl pH 7.9, 2 mM EDTA, 125 mM NaCl, 0.05% SDS, 1% Triton X-100, 1X complete protease inhibitor cocktail]; (2) high salt [20 mM Tris-HCl pH 7.9, 2 mM EDTA, 500 mM NaCl, 0.05% SDS, 1% Triton X-100, 1X complete protease inhibitor cocktail]; (3) LiCl [20 mM Tris-HCl pH 7.9, 1 mM EDTA, 250 mM LiCl, 1% NP-40, 1% sodium deoxycholate, 1X complete protease inhibitor cocktail]; (4) 1X Tris-EDTA [TE] containing 1X complete protease inhibitor cocktail. The precipitated immune complexes were transferred to a new tube in 1X TE before incubation in Elution Buffer (100 mM NaHCO3, 1% SDS) overnight at 65°C to de-crosslink and elute. Eluates were treated with DNase-free RNase (Roche, 11119915001) for 30 mins at 37°C, followed by 50 μg proteinase K (Life Technologies, 2542) for 2 hours at 55°C. Genomic DNA was purified by phenol chloroform extraction as previously described (27). ChIPed DNA was analyzed by qPCR. Non-specific background signals were determined using rabbit IgG isotype ChIP. The data were expressed as percent input.

### ChIP-qPCR primers

The following primer sets were used for ChIP-qPCR.

- *P2RY2* enh. forward: 5’-AGGTAAGAGCTGAGGAGCCC-3’
- *P2RY2* enh. reverse: 5’-CTTGACACCCCCAAGTGAGT-3’
- *NRIP1* enh. forward: 5’-GGCAGGTGATGATAACTCTGTAAGAAACC-3’
- *NRIP1* enh. reverse: 5’-GACAAACTGAACTTCTATACAGAGCTGCT-3’
- *TFF1* enh. forward: 5’-GATTTCCAACAAGATCTGCAACCACAGGGACG-3’
- *TFF1* enh. reverse: 5’-CTGACCCTCTGAAGCTCAAGGGCAGCTGGTGA-3’
- *MBOAT1* enh. forward: 5’-CTTCCAAACTCGCAAGCCAC-3’
- *MBOAT1* enh. reverse: 5’-CTCCAGCAGGAGTGAGTGTG-3’

### RNA-sequencing

RNA-seq libraries were generated as previously described with the following modifications (28). Total RNA from MCF-7 cells treated as indicated was isolated using the Qiagen RNeasy Plus Mini kit (Qiagen, 74136) according to the manufacturer’s protocol. Total RNA was then enriched for mRNA using Dynabeads Oligo(dT)25 (Invitrogen, 61002). Further fragmentation, first-strand cDNA synthesis, and subsequent procedures have been done as described (29).The libraries were subjected to quality control analyses, such as optimizing the number of PCR cycles required to amplify each set of libraries, quantifying the final library yield by Qubit fluorometer (Invitrogen, Q32851), and checking the size distribution of the final library by DNA ScreenTape system (Agilent Technologies, G2964AA, 5067–5584, and 5067–5603). The libraries were barcoded with indices as described for the Illumina TruSeq RNA Prep kit, multiplexed, and sequenced using Illumina NextSeq (2000) by 100 bp single-end sequencing. The RNA-seq data can be accessed from NCBI GEO using the accession number GSE267914.

### Analysis of RNA-seq data

The following steps were used to analyze the RNA-seq data: (1) Raw data were analyzed using FastQC tool for quality control; (2) Paired-end total RNA-seq reads were aligned to the human reference genome (GRCh38/hg38) using Star with default parameters; (3) Output files were then converted into BED files using SAMtools and BEDtools; and (4) bigWig files for visualization were generated using BEDtools. Heatmaps were generated using Java TreeView (30,31). Analysis of gene ontology (GO) was performed using the DAVID (Database for Annotation, Visualization, and Integrated Discovery) tool (32).

### Enhancer motif analysis

Nearest neighboring enhancers were determined for each differentially upregulated gene as determined from RNA-seq by defining ERα enhancers based on previously published ChIP-seq data (18). Whenever a gene had multiple nearby enhancers, the closest ERα peak was used. Motif analyses were then performed on a ±200 bp window around the summit of the ERα binding sites using HOMER gene-based analysis and ranked based on p-values (33).

### GTEx and TCGA tissue expression analyses

The expression of *GRAMD1B* mRNA in normal and cancer tissues was determined based on RPKM values using GEPIA (34).

### Immunocytochemistry staining of patient tumor microarrays

A paraffin-embedded breast cancer patient tumor microarray was obtained from US Biomax (BR2082b). Deparaffinization was performed by incubation in 100% xylene (Fisher, X3P) twice for 5 mins at room temperature. Slides were then rehydrated with sequential incubation in 100%, 95%, 70%, and 50% (Pharmco, 111000200) for 3 mins each at room temperature. Antigen retrieval was performed by incubation in boiling citrate buffer (10mM sodium citrate pH 6.0, 0.05% Tween-20) for 20 mins. Slides were then washed twice in 1X PBS for 10 mins at room temperature. Slides were blocked for 1 hour at room temperature using blocking solution described above, and stained for ASTER-B using a rabbit polyclonal antibody diluted 1:200 in 1X PBS for 2 hours at room temperature. Slides were then washed three times in 1X PBS, then incubated with Alexa Fluor 488 goat anti-rabbit IgG (ThermoFisher, A-11008; RRID:AB_143165) at a 1:500 dilution in 1X PBS for 1 hour at room temperature. Finally, coverslips were placed on cells coated with VectaShield Antifade Mounting Medium with DAPI (Vector Laboratories, H-1200; RRID:AB2336790).

### Quantification and statistical analyses

All sequencing-based genomic experiments were performed a minimum of two times with independent biological samples. Statistical analyses for the genomic experiments were performed using standard genomic statistical tests as described above. All gene specific qPCR-based experiments were performed a minimum of three times with independent biological samples. Statistical analyses were performed using GraphPad Prism 9. All tests and p-values are provided in the corresponding figures or figure legends.

### Data availability statement

All data generated in this study are available upon request from the corresponding author. The RNA-seq data can be accessed from NCBI GEO using the accession number GSE267914.

## Author Contributions

H.B.K.: Conceptualization, formal analysis, methodology, investigation, visualization, validation, writing-original draft. W.L.K.: Conceptualization, project development, funding acquisition, supervision, methodology, validation, project administration, visualization, writing-final draft.

## Acknowledgements

We would like to acknowledge Tulip Nandu for help in the computational analysis of RNA-seq data, Cristel Camacho for critical review of the manuscript, and Mikayla Stokes for scientific discussions. This work was supported by grants from the NIH/National Institute of Diabetes and Digestive and Kidney Diseases (NIH/NIDDK; R01 DK069710) and funds from the Cecil H. and Ida Green Center for Reproductive Biology Sciences Endowment to W.L.K.

## Note

This article contains no supplementary data.

## Supplementary Figure Legends

**Supplementary Figure S1. Accumulation of cellular E2 under low dose or prolonged treatment of MCF-7 cells**

**(A)** Measurement of intracellular E2 levels in MCF-7 cells by ELISA-based quantification upon continuous or without conditions with 100 pM E2. Initial treatment with E2 was 40 min, followed by 20 min of washout.

**(B)** Cellular E2 levels in MCF-7 cells upon continuous treatment with 100 nM E2 (left) or 100 pM (right) for up to 180 minutes as measured by ELISA-based quantification.

**(C)** Cellular E2 levels in MCF-7 cells upon acute treatment with 100 nM E2 for 40 min followed by prolonged washout with steroid-free medium for up to 24 hours.

**(D)** Gene ontology analysis for upregulated genes in continuous and washout groups. The top 5 terms as ranked by p-value (*top*) and selected terms of interest (*bottom*) are shown.

**Supplementary Figure S2. Motif analysis of nearest neighboring enhancers by bin.**

**(A)** Table showing number of significantly upregulated genes in either continuous or washout groups found in each bin.

**(B)** Schematic indicating that nearest neighboring enhancer analyses with ERα-ChIP seq data were used for subsequent motif analysis.

**(C)** Motifs of nearest at neighboring enhancers to genes analyzed by bin using HOMER. The top 5 motifs in each bin are shown as ranked by p-values in the continuous treatment group. Dashed boxes highlight motifs that are no longer significant in the washout group.

**Supplementary Figure S3. Modulation of ASTER-B or ERα expression effects E2 cellular retention.**

**(A)** Washout with hormone-free stripped medium after acute treatment with 100 nM E2-Glow reagent for scramble siRNA (*top*) or *GRAMD1B* knockdown (*bottom*) in MCF-7 cells. Initial treatment with E2 was 40 min, followed by 20 min of washout. Quantification of relative intensity of washout experiment images from n = 2 independent biological replicates normalized to values at t = 0 min (time of washout).

**(B)** Western blot showing knockdown of ASTER-B and ERα in MCF-7 cells.

**(C)** Measurement of intracellular E2 levels by ELISA-based quantification upon ASTER-B or ERα depletion in MCF-7 cells.

**(D)** Western blot showing ectopic expression of ERα-FLAG in HEK-293T cells.

**(E)** Measurement of intracellular E2 levels by ELISA-based quantification upon ectopic overexpression of ERα in HEK-293T cells. Initial treatment with E2 was 40 min, followed by 20 min of washout.

**(F)** Western blot of ASTER-B and ERα expression upon 100 nM E2 treatment over 6 hours in MCF-7 cells.

**Supplementary Figure S4. Regulation of gene expression is modulated by ASTER-B expression.**

Molecular assays showing the effect of *GRAMD1B* mRNA knockdown on E2-dependent genes expression outcomes.

**(A)** Chromatin occupancy of ERα at enhancer regions of E2-regulated genes (*TFF1* and *MBOAT1*) determined by ERα ChIP-qPCR. Initial treatment with E2 was 40 min, followed by 20 min of washout.

**(B)** Expression at E2-regulated genes (*TFF1* and *MBOAT1*) determined by RT-qPCR.

For statistical analysis, comparisons were made versus the vehicle control for each treatment group. Each bar represents the mean + range for n = 2. Initial treatment with E2 was 40 min, followed by 1 hour of washout. Significance was determined using multiple Student’s unpaired t-tests (non-parametric) with a Šidák-Bonferroni correction. Significance values were assigned as follows: * p<0.033, ** p<0.002, *** p<0.0001, **** p<0.00001.

**Supplementary Figure S5. Luminal B breast cancer and tissue specific gene expression of GRAMD1B.**

**(A)** Kaplan-Meier survival plots of luminal B subtype breast cancer patients with high (*red line*) or low (*blue line*) *GRAMD1B* mRNA expression level over a 5-year period. P-values were calculated using the Log-rank test.

**(B)** Tables showing breast cancer stage (*top*), histological subtype (*middle*), and receptor status (*bottom*) of BC tumor microarray categorized by ASTER-B localization.

**(C)** Tissue specific expression of *GRAMD1B* mRNA in humans from genome tissue expression (GTEx).

## References

1. Hanker AB, Sudhan DR, Arteaga CL. Overcoming Endocrine Resistance in Breast Cancer. Cancer Cell 2020;37(4):496–513 doi 10.1016/j.ccell.2020.03.009.

2. Endogenous H, Breast Cancer Collaborative G, Key TJ, Appleby PN, Reeves GK, Travis RC, et al. Sex hormones and risk of breast cancer in premenopausal women: a collaborative reanalysis of individual participant data from seven prospective studies. Lancet Oncol 2013;14(10):1009–19 doi 10.1016/S1470-2045(13)70301-2.

3. Zhang X, Tworoger SS, Eliassen AH, Hankinson SE. Postmenopausal plasma sex hormone levels and breast cancer risk over 20 years of follow-up. Breast Cancer Res Treat 2013;137(3):883–92 doi 10.1007/s10549-012-2391-z.

4. Pan H, Gray R, Braybrooke J, Davies C, Taylor C, McGale P, et al. 20-Year Risks of Breast-Cancer Recurrence after Stopping Endocrine Therapy at 5 Years. N Engl J Med 2017;377(19):1836–46 doi 10.1056/NEJMoa1701830.

5. Nelson LR, Bulun SE. Estrogen production and action. J Am Acad Dermatol 2001;45(3 Suppl):S116–24 doi 10.1067/mjd.2001.117432.

6. Chumsri S, Howes T, Bao T, Sabnis G, Brodie A. Aromatase, aromatase inhibitors, and breast cancer. J Steroid Biochem Mol Biol 2011;125(1-2):13–22 doi 10.1016/j.jsbmb.2011.02.001.

7. Cuzick J, Sestak I, Baum M, Buzdar A, Howell A, Dowsett M, et al. Effect of anastrozole and tamoxifen as adjuvant treatment for early-stage breast cancer: 10-year analysis of the ATAC trial. Lancet Oncol 2010;11(12):1135–41 doi 10.1016/S1470-2045(10)70257-6.

8. Morden JP, Alvarez I, Bertelli G, Coates AS, Coleman R, Fallowfield L, et al. Long-Term Follow-Up of the Intergroup Exemestane Study. J Clin Oncol 2017;35(22):2507–14 doi 10.1200/JCO.2016.70.5640.

9. Carlson RW, Hudis CA, Pritchard KI, National Comprehensive Cancer Network Breast Cancer Clinical Practice Guidelines in O, American Society of Clinical Oncology Technology Assessment on the Use of Aromatase I, St Gallen International Expert Consensus on the Primary Therapy of Early Breast C. Adjuvant endocrine therapy in hormone receptor-positive postmenopausal breast cancer: evolution of NCCN, ASCO, and St Gallen recommendations. J Natl Compr Canc Netw 2006;4(10):971–9 doi 10.6004/jnccn.2006.0082.

10. Fortunati N, Catalano MG, Boccuzzi G, Frairia R. Sex Hormone-Binding Globulin (SHBG), estradiol and breast cancer. Mol Cell Endocrinol 2010;316(1):86–92 doi 10.1016/j.mce.2009.09.012.

11. Haynes BP, Straume AH, Geisler J, A’Hern R, Helle H, Smith IE, et al. Intratumoral estrogen disposition in breast cancer. Clin Cancer Res 2010;16(6):1790–801 doi 10.1158/1078-0432.CCR-09-2481.

12. Takagi K, Ishida T, Miki Y, Hirakawa H, Kakugawa Y, Amano G, et al. Intratumoral concentration of estrogens and clinicopathological changes in ductal carcinoma in situ following aromatase inhibitor letrozole treatment. Br J Cancer 2013;109(1):100–8 doi 10.1038/bjc.2013.284.

13. van Landeghem AA, Poortman J, Nabuurs M, Thijssen JH. Endogenous concentration and subcellular distribution of estrogens in normal and malignant human breast tissue. Cancer Res 1985;45(6):2900–6.

14. Suzuki T, Miki Y, Ohuchi N, Sasano H. Intratumoral estrogen production in breast carcinoma: significance of aromatase. Breast Cancer 2008;15(4):270–7 doi 10.1007/s12282-008-0062-z.

15. Takagi M, Miki Y, Miyashita M, Hata S, Yoda T, Hirakawa H, et al. Intratumoral estrogen production and actions in luminal A type invasive lobular and ductal carcinomas. Breast Cancer Res Treat 2016;156(1):45–55 doi 10.1007/s10549-016-3739-6.

16. Xiao X, Kennelly JP, Feng AC, Cheng L, Romartinez-Alonso B, Bedard A, et al. Aster-B-dependent estradiol synthesis protects female mice from diet-induced obesity. J Clin Invest 2024;134(4) doi 10.1172/JCI173002.

17. Sandhu J, Li S, Fairall L, Pfisterer SG, Gurnett JE, Xiao X, et al. Aster Proteins Facilitate Nonvesicular Plasma Membrane to ER Cholesterol Transport in Mammalian Cells. Cell 2018;175(2):514–29 e20 doi 10.1016/j.cell.2018.08.033.

18. Franco HL, Nagari A, Kraus WL. TNFalpha signaling exposes latent estrogen receptor binding sites to alter the breast cancer cell transcriptome. Mol Cell 2015;58(1):21–34 doi 10.1016/j.molcel.2015.02.001.

19. Khanna P, Chua PJ, Wong BSE, Yin C, Thike AA, Wan WK, et al. GRAM domain-containing protein 1B (GRAMD1B), a novel component of the JAK/STAT signaling pathway, functions in gastric carcinogenesis. Oncotarget 2017;8(70):115370–83 doi 10.18632/oncotarget.23265.

20. Nguyen Ba AN, Pogoutse A, Provart N, Moses AM. NLStradamus: a simple Hidden Markov Model for nuclear localization signal prediction. BMC Bioinformatics 2009;10:202 doi 10.1186/1471-2105-10-202.

21. Laraia L, Friese A, Corkery DP, Konstantinidis G, Erwin N, Hofer W, et al. The cholesterol transfer protein GRAMD1A regulates autophagosome biogenesis. Nat Chem Biol 2019;15(7):710–20 doi 10.1038/s41589-019-0307-5.

22. Cancer Genome Atlas Research N. Comprehensive genomic characterization defines human glioblastoma genes and core pathways. Nature 2008;455(7216):1061–8 doi 10.1038/nature07385.

23. Murase K, Yanai A, Saito M, Imamura M, Miyagawa Y, Takatsuka Y, et al. Biological characteristics of luminal subtypes in pre- and postmenopausal estrogen receptor-positive and HER2-negative breast cancers. Breast Cancer 2014;21(1):52–7 doi 10.1007/s12282-012-0348-z.

24. Kraus WL, Kadonaga JT. p300 and estrogen receptor cooperatively activate transcription via differential enhancement of initiation and reinitiation. Genes Dev 1998;12(3):331–42 doi 10.1101/gad.12.3.331.

25. Kraus WL, Montano MM, Katzenellenbogen BS. Identification of multiple, widely spaced estrogen-responsive regions in the rat progesterone receptor gene. Mol Endocrinol 1994;8(8):952–69 doi 10.1210/mend.8.8.7997237.

26. Murakami S, Nagari A, Kraus WL. Dynamic assembly and activation of estrogen receptor alpha enhancers through coregulator switching. Genes Dev 2017;31(15):1535–48 doi 10.1101/gad.302182.117.

27. Murakami S, Li R, Nagari A, Chae M, Camacho CV, Kraus WL. Distinct Roles for BET Family Members in Estrogen Receptor alpha Enhancer Function and Gene Regulation in Breast Cancer Cells. Mol Cancer Res 2019;17(12):2356–68 doi 10.1158/1541-7786.MCR-19-0393.

28. Hou TY, Kraus WL. Analysis of estrogen-regulated enhancer RNAs identifies a functional motif required for enhancer assembly and gene expression. Cell Rep 2022;39(11):110944 doi 10.1016/j.celrep.2022.110944.

29. Zhong S, Joung JG, Zheng Y, Chen YR, Liu B, Shao Y, et al. High-throughput illumina strand-specific RNA sequencing library preparation. Cold Spring Harb Protoc 2011;2011(8):940–9 doi 10.1101/pdb.prot5652.

30. Saldanha AJ. Java Treeview--extensible visualization of microarray data. Bioinformatics 2004;20(17):3246–8 doi 10.1093/bioinformatics/bth349.

31. Dobin A, Davis CA, Schlesinger F, Drenkow J, Zaleski C, Jha S, et al. STAR: ultrafast universal RNA-seq aligner. Bioinformatics 2013;29(1):15–21 doi 10.1093/bioinformatics/bts635.

32. Huang DW, Sherman BT, Lempicki RA. Systematic and integrative analysis of large gene lists using DAVID bioinformatics resources. Nat Protoc 2009;4(1):44–57 doi 10.1038/nprot.2008.211.

33. Heinz S, Benner C, Spann N, Bertolino E, Lin YC, Laslo P, et al. Simple combinations of lineage-determining transcription factors prime cis-regulatory elements required for macrophage and B cell identities. Mol Cell 2010;38(4):576–89 doi 10.1016/j.molcel.2010.05.004.

34. Tang Z, Li C, Kang B, Gao G, Li C, Zhang Z. GEPIA: a web server for cancer and normal gene expression profiling and interactive analyses. Nucleic Acids Res 2017;45(W1):W98–W102 doi 10.1093/nar/gkx247.

