## Supplementary Figures S1-S5 + legends for "Intracellular Retention of Estradiol is Mediated by GRAM Domain-Containing Protein ASTER-B in Breast Cancer Cells"

This document contains:
• Supplementary Figures S1-S5
• Legends for Supplementary Figures S1-S5

### SUPPLEMENTARY FIGURES

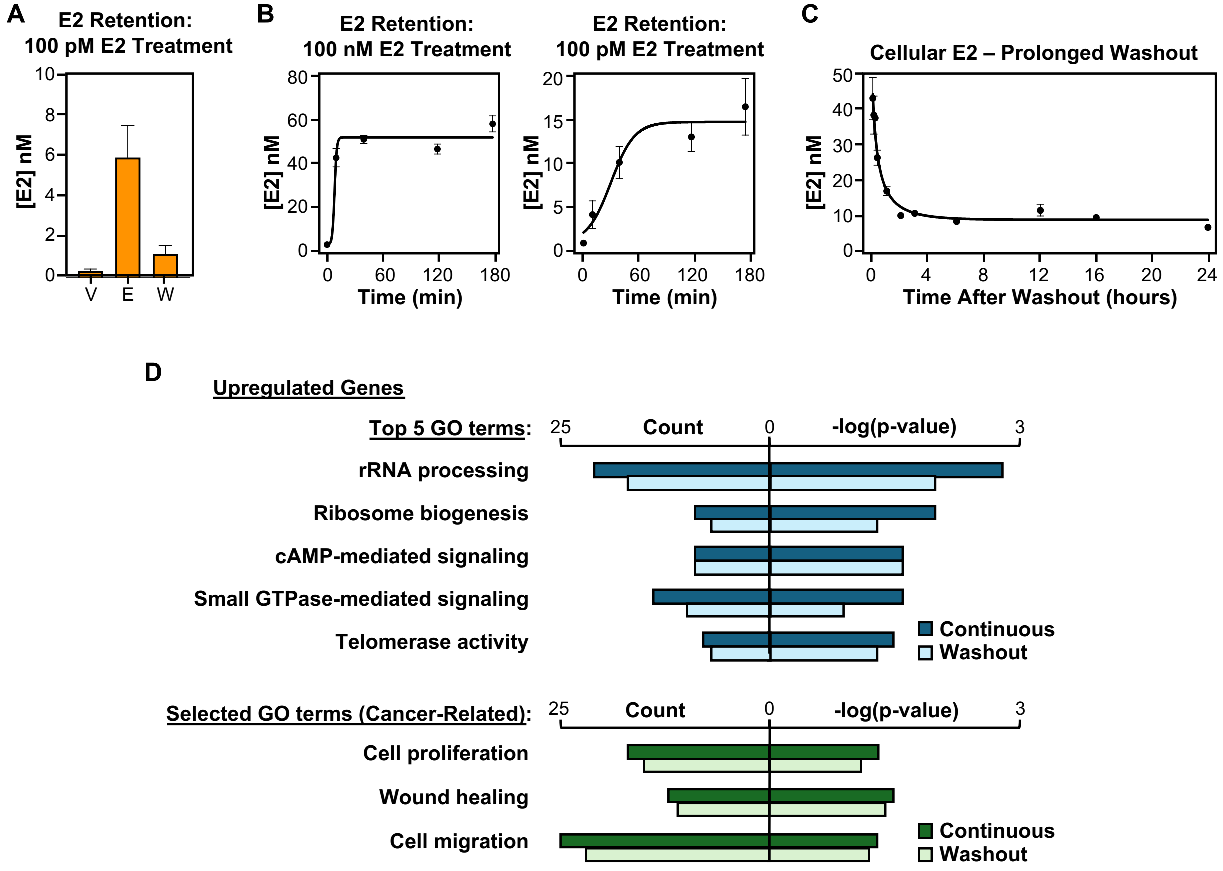


**Supplementary Figure S1. Accumulation of cellular E2 under low dose or prolonged treatment of MCF-7 cells**

**(A)** Measurement of intracellular E2 levels in MCF-7 cells by ELISA-based quantification upon continuous or without conditions with 100 pM E2. Initial treatment with E2 was 40 min, followed by 20 min of washout.

**(B)** Cellular E2 levels in MCF-7 cells upon continuous treatment with 100 nM E2 (left) or 100 pM (right) for up to 180 minutes as measured by ELISA-based quantification.

**(C)** Cellular E2 levels in MCF-7 cells upon acute treatment with 100 nM E2 for 40 min followed by prolonged washout with steroid-free medium for up to 24 hours.

**(D)** Gene ontology analysis for upregulated genes in continuous and washout groups. The top 5 terms as ranked by p-value (*top*) and selected terms of interest (*bottom*) are shown.

**A**

| Bin | Number of Genes (Continuous) | Number of Genes (Washout) |
|---|---|---|
| 1 | 16 | 18 |
| 2 | 174 | 91 |
| 3 | 436 | 281 |
| 4 | 1,135 | 969 |
| 5 | 1,210 | 901 |

**B**

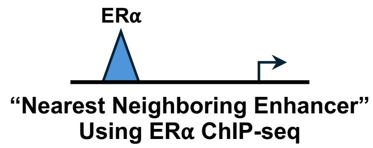


**C**

| Bin 2: Motif | Logo | p-value Control | p-value Washout |
|---|---|---|---|
| ERE | | $1 \times 10^{-280}$ | $1 \times 10^{-250}$ |
| THRb | | $1 \times 10^{-150}$ | $1 \times 10^{-40}$ |
| FOXM1 | | $1 \times 10^{-40}$ | |
| RARA | | $1 \times 10^{-36}$ | |
| GATA6 | | $1 \times 10^{-35}$ | |

| Bin 3: Motif | Logo | p-value Control | p-value Washout |
|---|---|---|---|
| ERE | | $1 \times 10^{-300}$ | $1 \times 10^{-190}$ |
| GATA3 | | $1 \times 10^{-380}$ | $1 \times 10^{-30}$ |
| FOXA1 | | $1 \times 10^{-370}$ | |
| ARID5a | | $1 \times 10^{-360}$ | |
| FOXA2 | | $1 \times 10^{-350}$ | |

| Bin 4: Motif | Logo | p-value Control | p-value Washout |
|---|---|---|---|
| ERE | | $1 \times 10^{-410}$ | $1 \times 10^{-300}$ |
| THRb | | $1 \times 10^{-130}$ | $1 \times 10^{-88}$ |
| MAFG | | $1 \times 10^{-630}$ | |
| GATA3 | | $1 \times 10^{-570}$ | |
| FOXA1 | | $1 \times 10^{-39}$ | $1 \times 10^{-30}$ |

| Bin 5: Motif | Logo | p-value Control | p-value Washout |
|---|---|---|---|
| ERE | | $1 \times 10^{-360}$ | $1 \times 10^{-280}$ |
| RARA | | $1 \times 10^{-78}$ | |
| MAFG | | $1 \times 10^{-55}$ | |
| GATA3 | | $1 \times 10^{-38}$ | |
| FOXA1 | | $1 \times 10^{-37}$ | $1 \times 10^{-38}$ |

**Supplementary Figure S2. Motif analysis of nearest neighboring enhancers by bin.**
**(A)** Table showing number of significantly upregulated genes in either continuous or washout groups found in each bin.
**(B)** Schematic indicating that nearest neighboring enhancer analyses with ERα-ChIP seq data were used for subsequent motif analysis.
**(C)** Motifs of nearest at neighboring enhancers to genes analyzed by bin using HOMER. The top 5 motifs in each bin are shown as ranked by p-values in the continuous treatment group. Dashed boxes highlight motifs that are no longer significant in the washout group.

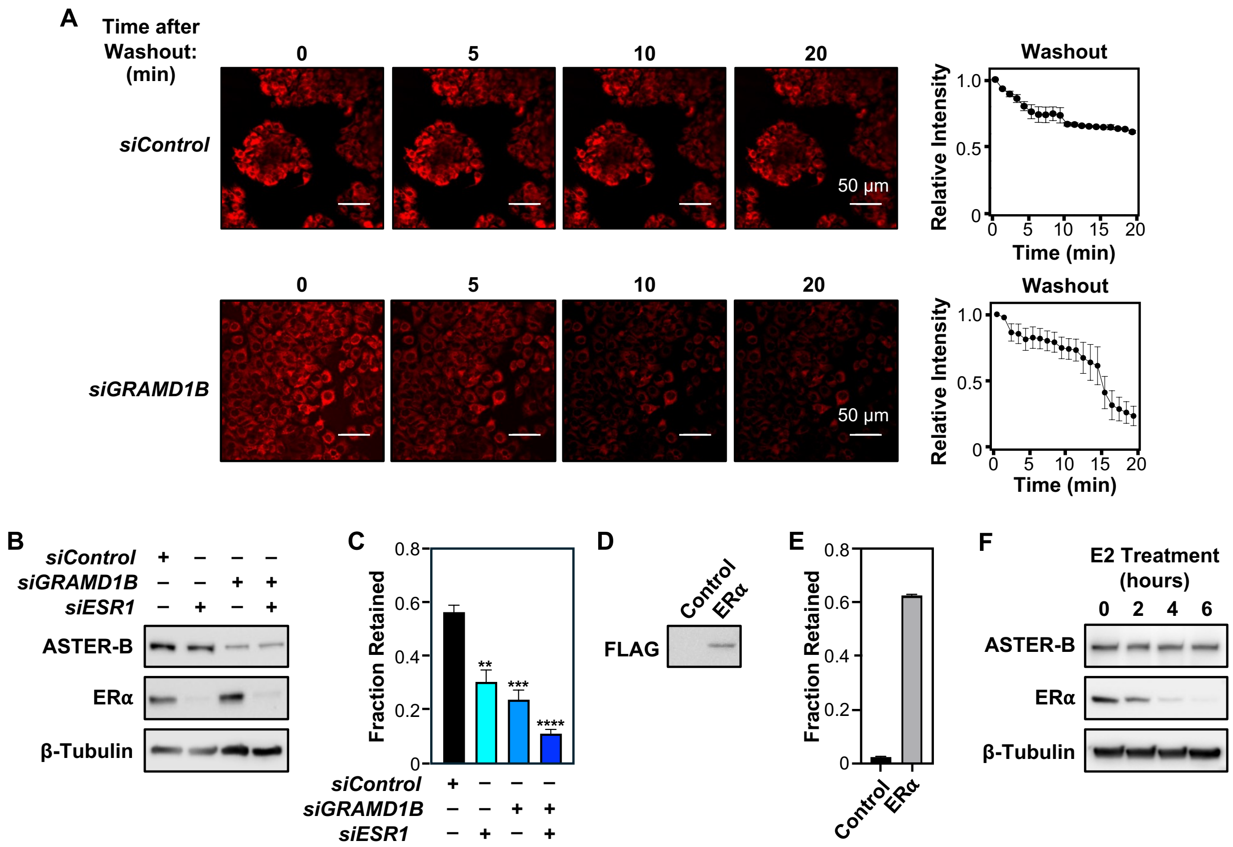


**Supplementary Figure S3. Modulation of ASTER-B or ERα expression effects E2 cellular retention.**

**(A)** Washout with hormone-free stripped medium after acute treatment with 100 nM E2-Glow reagent for scramble siRNA (*top*) or *GRAMD1B* knockdown (*bottom*) in MCF-7 cells. Initial treatment with E2 was 40 min, followed by 20 min of washout. Quantification of relative intensity of washout experiment images from n = 2 independent biological replicates normalized to values at t = 0 min (time of washout).

**(B)** Western blot showing knockdown of ASTER-B and ERα in MCF-7 cells.

**(C)** Measurement of intracellular E2 levels by ELISA-based quantification upon ASTER-B or ERα depletion in MCF-7 cells.

**(D)** Western blot showing ectopic expression of ERα-FLAG in HEK-293T cells.

**(E)** Measurement of intracellular E2 levels by ELISA-based quantification upon ectopic overexpression of ERα in HEK-293T cells. Initial treatment with E2 was 40 min, followed by 20 min of washout.

**(F)** Western blot of ASTER-B and ERα expression upon 100 nM E2 treatment over 6 hours in MCF-7 cells.

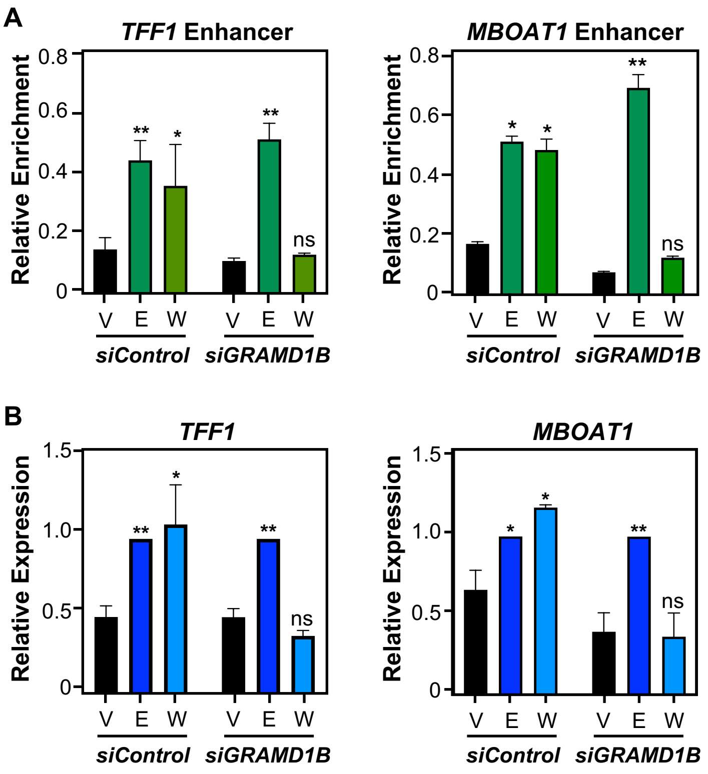


**Supplementary Figure S4. Regulation of gene expression is modulated by ASTER-B expression.**

Molecular assays showing the effect of GRAMD1B mRNA knockdown on E2-dependent genes expression outcomes.

**(A)** Chromatin occupancy of ERα at enhancer regions of E2-regulated genes (*TFF1* and *MBOAT1*) determined by ERα ChIP-qPCR. Initial treatment with E2 was 40 min, followed by 20 min of washout.

**(B)** Expression at E2-regulated genes (*TFF1* and *MBOAT1*) determined by RT-qPCR.

For statistical analysis, comparisons were made versus the vehicle control for each treatment group. Each bar represents the mean + range for n = 2. Initial treatment with E2 was 40 min, followed by 1 hour of washout. Significance was determined using multiple Student's unpaired t-tests (non-parametric) with a Šidák-Bonferroni correction. Significance values were assigned as follows: * $p<0.033$, ** $p<0.002$, *** $p<0.0001$, **** $p<0.00001$.

A

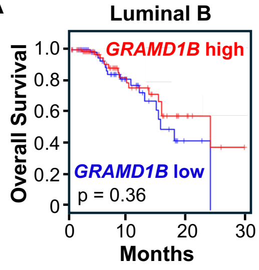


B

| BC Stage | Nuclear ASTER-B | Cytoplasmic ASTER-B |
|---|---|---|
| None | 8 | 19 |
| I | 2 | 5 |
| II | 9 | 24 |
| III | 5 | 2 |

| Histological Subtype | Nuclear ASTER-B | Cytoplasmic ASTER-B |
|---|---|---|
| Metastatic | 2 | 26 |
| Invasive Ductal Carcinoma | 15 | 15 |
| Invasive Lobular Carcinoma | 2 | 1 |
| Intraductal Carcinoma | 3 | 5 |
| Normal Breast Tissue | 2 | 0 |
| Squamous Cell Carcinoma | 0 | 1 |
| Atypical Hyperplasia | 0 | 1 |
| Plasma Cell Mastitis | 0 | 1 |

| Receptor Status | Nuclear ASTER-B | Cytoplasmic ASTER-B |
|---|---|---|
| ER+ | 19 | 26 |
| PGR+ | 14 | 17 |
| HER2+ | 9 | 20 |
| None | 0 | 8 |

C

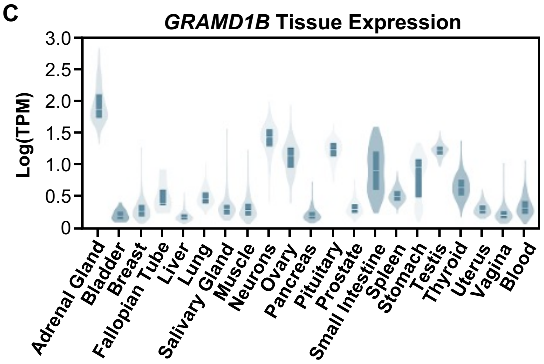


**Supplementary Figure S5. Luminal B breast cancer and tissue specific gene expression of GRAMD1B.**

**(A)** Kaplan-Meier survival plots of luminal B subtype breast cancer patients with high (*red line*) or low (*blue line*) *GRAMD1B* mRNA expression level over a 5-year period. P-values were calculated using the Log-rank test.

**(B)** Tables showing breast cancer stage (*top*), histological subtype (*middle*), and receptor status (*bottom*) of BC tumor microarray categorized by ASTER-B localization.

**(C)** Tissue specific expression of *GRAMD1B* mRNA in humans from genome tissue expression (GTEx).
